# Macrophage Immune-Competent Colon Assembloids for Functional Interrogation of Neuroinflammation-Induced Colonic Dysmotility

**DOI:** 10.1101/2025.08.07.669171

**Authors:** Claudia A. Collier, Karla I. Ortega Sandoval, Aelita Salikhova, Shrinarayanee Rengarajan, Anvitha Tharakesh, Alana Aristimuno Millan, Shanthi Srinivasan, Shreya A. Raghavan

**Affiliations:** Department of Biomedical Engineering, Texas A&M University; Department of Medicine, Division of Digestive Diseases, Emory University

## Abstract

Functional gastrointestinal disorders (FGIDs) affect ∼40% of the global population and are frequently characterized by colonic dysmotility. Symptomatic manifestations of colonic dysmotility significantly reduce quality of life in inflammatory bowel disease (IBD), diabetes, and Gulf War Illness (GWI). Current in vitro models lack the integration of functional physiology with immune and neuronal complexity required to establish causal links between neuroinflammation and dysmotility. Here, an immune-competent bioengineered colon assembloid is introduced that integrates multiple cell types of the external colonic wall, along with functional readouts of motility. Within bioengineered colon assembloids, various inflammatory insults resulted in enteric neuroinflammation, cascading to changes in colonic motility. Key mechanisms of dysmotility following inflammatory insult within the bioengineered colon assembloids included impaired neuronal regeneration, and aberrant smooth muscle remodeling. The bioengineered colon assembloid model mimicked diverse aspects of enteric neuroinflammation. Ultimately, the platform offers a physiologically relevant avenue to interrogate neuroimmune crosstalk and dissect mechanisms of colonic dysmotility, paving the way to new therapeutic strategies to improve colonic motility.

## 1. INTRODUCTION

Functional gastrointestinal disorders (FGIDs) are a significant global health burden^1^, affecting an estimated 40% of people globally^2^. FGIDs occur in individuals with Inflammatory Bowel Disease (IBD)^3^, Gulf War Illness (GWI)^4, 5^, and even secondary to diabetes^6^. At the heart of FGIDs is altered colonic motility (colonic dysmotility). Yet, the mechanistic underpinnings of FGID-related colonic dysmotility are still poorly understood, and consequently, our ability to design targeted therapeutics to curtail it lag. This gap is partly due to the lack of nuanced in vitro model systems that can functionally integrate multi-cellular communication with motility-focused read outs.

The smooth muscle layers of the colonic muscularis externa integrate input from the enteric nervous system (ENS), tissue-resident immune cells like macrophages, and other interstitial cells to produce colonic motility^7^. Coordinated colonic motility occurs via reciprocal interactions between many of these individual cell types, with a beautifully complex regulatory network^7^. Chief among these are neuro-immune interactions, well studied in infection, inflammation, and mucosal barrier function^8,9^. From whole animal models, we deeply appreciate the evolving role of neuro-immune interactions in colonic motility, dysmotility and FGIDs^10,11^, especially those between macrophages and enteric nerves^12^.

Subtle alterations in the connectivity and biochemistry of the ENS impact colonic motility patterns^13^. Tolerogenic macrophages support the regeneration of damaged enteric nerves from a tissue-resident enteric neuronal progenitor pool, by clearing neuronal debris^14^. Activated pro-inflammatory macrophages release cytokines that impair neuronal regeneration and smooth muscle contraction^15, 16^, thereby affecting colonic motility. Collectively, the presence of altered neuroplasticity, increased pro-inflammatory cytokines and altered colonic motility supports a model in which macrophages, smooth muscle cells of the colon, and enteric neurons interact to contribute to the symptoms of colonic dysmotility^12, 17, 18^.

These foundational insights have been obtained from experimental mouse models, linking functional physiology to histological observations. A hallmark study demonstrated that macrophage-ENS interactions directed colonic motility via bone morphogenetic protein-2 (BMP-2) secretion^12^. Another experimental mouse model correlated the role of macrophages in myenteric plexus repair to intestinal dysmotility following surgical trauma^17^. Our own research with mouse models of Gulf War Illness demonstrated that toxic exposures reprogrammed macrophages and induced enteric neuronal progenitor cell apoptosis, leading to chronically altered colonic motility^19, 20^. Similar disruptions in neuroplasticity and altered colonic motility are routinely reported in mouse models of inflammatory bowel disease and FGIDs, often accompanied by increased pro-inflammatory cytokines^11, 18, 21^. However, establishing signal specificity and interrogating molecular mechanisms directing colonic motility independent of the microbiome and other systemic factors is tedious and challenging in experimental animal models. Herein, bioengineering and in vitro modeling offer a fantastic opportunity for tight variable isolation for mechanistic studies^22, 23^.

Ongoing advances in organoid technologies^24^, discovery and use of postnatal and adult human enteric neural progenitor cells^25^, and gene editing have paved the way towards in vitro bioengineered systems capable of nuanced mechanistic dissection in enteric neuro-inflammation. Murine colonic organoids co-cultured with sensory neurons demonstrated a shared calcium influx pathway in macrophage-nerve interactions that may lead to pain in FGIDs^26^. Colonic organoids incorporating epithelial and immune components have been previously used to investigate immune and cytokine signaling in IBD^27^. Organoids, however, are largely epithelial-dominant systems with limited utility in studying neuro-immune crosstalk and motility. For example, co-culture of human intestinal organoids with induced pluripotent stem cell-derived neural progenitor cells showed neuroglial structures like the myenteric plexus. While this system demonstrated transient calcium waves, similar to neuronal activity that regulates colonic motility, it falls short of being able to quantify motility and contractility^28^. This is because organoids lack the active smooth muscle component of the outer layers of the colon that generate motility forces^29^.

Assembloid systems on the other hand offer a construction strategy that enables multi-cellular structures within directed architectures^30, 31^. Colon assembloid systems have been successfully used to study stromal cell sustenance of epithelial crypts^32^, and macrophage involvement in mucosal immunity^33^. Silk scaffold directed assembly of intestinal epithelium, myofibroblasts and differentiated enteric nerves enabled investigation of mucosal barrier function^34^. More recently, ENS-rich assembloids were used to study the dynamics and mechanisms of ENS regeneration and neural elongation during inflammation^35^. Despite these successes, current assembloid models also do not integrate functional readouts of colonic motility including spontaneous or evoked contractions in response to exogenous stimuli, thereby limiting their ability to capture how neuroinflammation drives colonic motility (and dysmotility).

The objective of the current study, therefore, was to create a bioengineered colon assembloid that integrated macrophage-ENS crosstalk with smooth muscle contractility. Our model guides the self-assembly of key cell types of the colonic wall that regulate colonic motility. These cell types include smooth muscle cells, macrophages, and an enteric neural circuit derived from enteric neuronal progenitor cells. This platform structurally mimics the outer colon layer and enables real-time assessment of tissue contractility. In this study, we show that bioengineered colon assembloids respond to inflammatory insults by altering macrophage activation, neural structure and composition, ultimately impacting colonic motility. Beyond its relevance to specific disease models, this system serves as a foundational platform for deeper mechanistic inquiry into neuroimmune processes driving colonic motility and dysmotility.

## 2. METHODS

### 2.1 Materials and reagents

All tissue culture media, and supplements were purchased from ThermoFisher Scientific (Waltham, MA, USA), unless otherwise specified. Growth factors and proteins were purchased from Peprotech (Cranbury, NJ, USA). All conjugated antibodies were purchased from Santa Cruz (Dallas, TX, USA), Cell Signaling Technology (Danvers, MA, USA), or Abcam (Cambridge, MA, USA), unless specified otherwise. Pyridostigmine bromide, Acetycholine (ACh), Tetrodotoxin (TTX) were purchased from Sigma Aldrich (St. Louis, MO, USA).

### 2.2 Cell lines and culture

#### 2.2.1 Isolation And Culture of Immortalized Colonic Smooth Muscle Cells (icSMCs)

Primary murine colonic smooth muscle cells were isolated from adult mouse colons aged 20-28 weeks. During dissection, the sigmoid colon was isolated via sharp dissection^19, 36, 37^. Dissected sigmoid colons were cleaned and prepped for enzymatic digestion. Briefly, dissected colonic tissue was washed in sterile phosphate buffered saline, minced with sharp scissors, and digested in type II collagenase. Digested cells were filtered serially through a 70 and 40 μm mesh to remove debris, and plated in complete Dulbecco’s Modified Eagle Medium (DMEM) supplemented with 10% fetal bovine serum + 1% penicillin and streptomycin. After initial expansion, cells were transduced with EF1α-SV40 large T antigen (SV40T) lentiviral particles (GeneCopoeia, puromycin-selectable; Rockville, MD, USA) to establish a stable immortalized smooth muscle cell line (icSMC). Successfully transduced cells were selected using puromycin and maintained at 37°C with media changes every 48 hours.

#### 2.2.2 Culture Of Immortalized Bone Marrow Derived Macrophages

Immortalized bone marrow derived macrophages^38^ (iBMDMs) were a gift from Dr. Philip West’s laboratory at Jackson Labs, Bar Harbor ME. Cells were maintained in complete DMEM supplemented with 10% fetal bovine serum + 1% antibiotic-antimycotic at 37°C with 5% CO_2_.

#### 2.2.3 Culture Of Immortalized Fetal Enteric Neural Cells (IM-FENs)

The immortalized murine fetal enteric neuron (IM-FEN) cell line used in this study was generated and characterized by the Srinivasan Laboratory at Emory University^39, 40^. Briefly, IM-FENs were derived from E13 H-2Kb-tsA58 immortomouse embryos. Neural precursors were isolated using magnetic bead immunoselection targeting the low-affinity NGF receptor (Ngfr-p75). IM-FENs were maintained in a modified N2 medium supplemented with 10% fetal bovine serum, 10 ng/mL glial cell line-derived neurotrophic factor (GDNF), 20 U/mL recombinant mouse interferon-γ, and 1% penicillin/streptomycin. Cells were cultured at 33°C in a humidified incubator with 5% CO₂ to preserve their progenitor state. To induce neuronal differentiation, cells were transitioned to a 1:1 ratio of conditioned media collected from icSMCs and neurobasal-A medium supplemented with B-27 (serum-free), 2 mmol/L L-glutamine, 1% heat-inactivated fetal bovine serum, 1% penicillin/streptomycin, and 10 ng/mL GDNF. Cultures were maintained at 39°C and 5% CO₂ for up to 7 days to promote differentiation into mature enteric neurons.

### 2.3 Bioengineered colon assembloid formation and characterization

#### 2.3.1 Assembly

Bioengineered colon assembloids were fabricated using a two-layer collagen hydrogel deposition technique to enable directed self-assembly within cell-laden hydrogels. Briefly, 35mm dishes were prepared by coating with a thin layer of non-adhesive polydimethyl siloxane (PDMS; Sylgard® 184 silicone elastomer kit, Denver, CO, USA). The coating also contained a central silicone post (3mm diameter) to define the hollow lumen. Following curing over 48hrs, 35mm culture dishes had a non-adherent surface with a defined central post. Around this central post, two sequential layers of collagen hydrogels (1 mg/ml type I collagen; rat tail; ibidi; Madison, WI, USA) were patterned one after the other.

The following cell types were uniformly suspended within the first base collagen gel layer: (i) 400,000 cells/mL IM-FENs; (ii) 1,500 cells/mL iBMDMs; and (iii) 200,000 cells/mL icSMCs. 300µl of cell-laden gel mixture was cast around the central post and allowed to gel at 37°C for 15 minutes.

A second cell-laden collagen gel suspension was prepared using 600,000 cells/mL iCSMCs only. Once gelation of the first base layer with all three cell types was complete, 300µL of cell-laden second layer was patterned around the first layer and the post. The assembled structures were allowed to gel for an additional 30 minutes at 37°C. Following gelation, 2 mL of iCSMC media was added to each dish, and constructs were incubated at 37°C to promote self-assembly, alignment and compaction. After 24 hours, media was supplemented with neuronal differentiation media for a final 1:1 ratio of icSMC media: neural differentiation media to support differentiation of enteric neuronal progenitor cells. Formation of bioengineered colon assembloids was monitored via daily photography, ensuring compaction and tight assembly around luminal post. Compaction of the gel area was measured and monitored over 7 days.

#### 2.3.2 Visualization of multi-cellular compartments within bioengineered colon assembloids using whole mount immunofluorescence

Bioengineered colon assembloids were fixed in 4% paraformaldehyde for 30 minutes at room temperature, followed by three washes in phosphate-buffered saline (PBS). Assembloids were washed in 0.1M glycine for 20 minutes at room temperature followed by permeabilizing and blocking for 2 hours in 0.2% Triton-X and 10% goat serum. Fluorophore-conjugated antibodies targeting lineage-specific markers (**Supplementary Table 1**) were diluted in blocking buffer and incubated overnight at 4°C. Following incubation with antibodies, samples were then washed thoroughly, mounted between coverslips and imaged using Olympus Fluoview FV3000 Confocal Laser Scanning Microscope (RRID:SCR_021637). Each experimental condition was replicated across multiple biological samples (n ≥ 5). Maximum intensity projections of the XY plane were visualized to demonstrate overall structural organization of cell types within the bioengineered colon assembloids.

### 2.4 Inflammatory Stimulation of Bioengineered Colon Assembloids

Bioengineered colon assembloids were established and allowed to fully form between Days 1-3. Once compaction was achieved, inflammatory insult was initiated by adding 100µM Pyridostigmine Bromide (PB) or 100ng/mL Tumor Necrosis Factor-alpha (TNF-α) into the culture media of assembloids for 24hrs. PB is a toxic exposure linked to neuroinflammation and FGIDs in Gulf War Illness^19, 20^, while TNF-α is a cytokine observed in many FGIDs^41, 42^. Inflammatory insult was initiated at Day 3 and removed on Day 4.

#### 2.4.1 Defining “acute” and “chronic” lingering effects of inflammation following one-time inflammatory insult

An “acute” immediate effect was assessed by collecting assembloids at Day 4 for downstream processing. Assembloids were also allowed to recover for 3 days following one-time inflammatory insult (Days 4-7), based on previously documented studies establishing that in vitro macrophage activation dynamics changed over 72hrs^43, 44^. Any lingering “chronic” effects of one-time inflammatory insult were therefore assessed following recovery, at Day 7. Time-matched untreated assembloids were maintained as controls.

### 2.5 Characterization of the bioengineered colon assembloids following inflammatory insult for acute and chronic effects

Following assembly (1-3 days), and inflammatory insult, colon assembloids were processed for downstream analysis to investigate acute (Day 4) and chronic effects (Day 7) of inflammation. Analyses focused on targeted gene expression analysis, unbiased RNA-Sequencing analysis, cytokine profiling and quantitative whole mount immunofluorescence.

#### 2.5.1 RNA extraction and gene expression analysis

RNA was extracted from cultured cells or assembloids using a RNeasy extraction kit (Qiagen, Hilden, Germany). Bioengineered colon assembloids were first flash frozen in liquid nitrogen, minced in cold TRIzol™ LS Reagent (Invitrogen, Carlsbad, CA, USA) and extracted using the same kit following manufacture’s protocol. RNA concentrations were measured using a Nanodrop 2000 (ThermoFisher Scientific, Carlsbad, CA, USA) spectrophotometer, transcribed to cDNA using the high-fidelity cDNA transcription kit (Life Technologies, Carlsbad, CA, USA), and q-PCR was carried out in the 96-well format using a QuantStudio 3(Applied Biosystems, Foster City, CA, USA). Gene expression differences were quantified using the 2ΔΔCT method, using *Gapdh* as the housekeeping control, and reported as fold changes compared to a control sample. qPCR experiments were run in triplicates with 5 independent samples. Primer sets used are detailed in **Supplementary Table 2**. Gene expression was normalized to time-matched control colon assembloids that received no inflammatory insult. Fold changes in gene expression were reported at acute (Day 4) or chronic (Day 7) time points due to PB or TNF-α treatment, compared to time-matched untreated controls.

#### 2.5.2 RNA Sequencing and analysis

RNA was isolated from PB or TNF-α treatment bioengineered colon assembloids, or control untreated assembloids at Day 4 (acute; n=4) and Day 7 (chronic; n=3). Library preparation with poly(A) selection and 150-bp paired-end sequencing on an Illumina HiSeq 2500 were performed by Azenta (South Plainfield, NJ, USA). Raw sequencing data in FASTQ format were generated by Azenta Life Sciences using an Illumina platform. Azenta performed initial quality assessment, adapter trimming, alignment, and gene count quantification. Specifically, adapter sequences and low-quality bases were trimmed using Trimmomatic v0.36, and trimmed reads were aligned to the *Mus musculus* GRCm38 reference genome (Ensembl) using STAR aligner v2.5.2b. Gene-level quantification was performed using featureCounts from the Subread package v1.5.2, generating a raw gene count matrix for each sample.

All subsequent computational and statistical analyses were conducted in-house using RStudio (2025.05.1+513) and R packages listed below. Differential gene expression analysis (DGE) was performed using the DESeq2 package. Genes with p-value < 0.05 and absolute log2 fold change > 1.25 were defined as differentially expressed genes (DEGs).

Volcano plots were generated using ggplot2 and dplyr, and 3D principal component analysis (PCA) was performed using plotly and dplyr to visualize sample clustering based on transcriptomic profiles. Venn diagrams were used to compare the number of upregulated and downregulated DEGs across conditions relative to controls. Gene ontology (GO) enrichment analysis was conducted using the EnRichGO, clusterProfiler, org.Mm.eg.db, AnnotationDbi, and tidyverse packages. Biological process (BP) terms were assessed for DEGs with log2FC > 1.25 and p-value < 0.05.

#### 2.5.3 Cytokine profiling within bioengineered colon assembloids

Bioengineered colon assembloids were maintained as untreated controls, or subject to inflammatory insult (PB or TNF-α) at Day 3 with removal at Day 4. “Acute” effects were evaluated at Day 4, while “chronic” lingering effects were evaluated following 3 days of recovery, at Day 7. Assembloids were flash frozen with liquid nitrogen and collected in a RIPA+HALT solution. Supernatants were clarified by centrifugation at 500× *g* for 5 minutes, sterile filtered, and stored at −80°C until analysis. Protein concentration was measured using a Pierce BCA assay (Thermo-Fisher) following manufacturer’s protocols. Cytokine levels were evaluated using bead-based multiplex immunoassays and done according to the manufacturer’s protocol (R&D Systems, Minneapolis, MN). Equal amounts of protein/sample were used per assay, across acute and chronic time points for each inflammatory insult, compared to untreated time-matched controls. Magnetic bead washes and intermediary steps were performed using a Bio-Rad Bio-Plex Pro II plate washer and final luminescence was read on a Bio-Rad Bio-Plex analyzer. Each experimental condition was replicated across multiple biological samples (n ≥ 3). The following cytokines were assayed: CCL2 (C-C motif chemokine ligand 2), IL-1β (Interleukin-1 beta), IL-10 (Interleukin-10), TIMP-1 (Tissue inhibitor of metalloproteinases-1), CCL11 (C-C motif chemokine ligand 11), β-NGF (beta-nerve growth factor), IL-6 (Interleukin-6).

#### 2.5.4 Evaluation of architectural changes within bioengineered colon assembloids with inflammatory insult via whole mount immunofluorescence and quantification

Assembloids were formed, exposed to inflammatory insult, and fixed and stained at “acute” Day 4 or “chronic” Day 7 time points. Time-matched untreated controls were also fixed and stained using methods outlined in Section *2.3.2*. Whole mount immunofluorescence was performed to visualize macrophages and their activation via co-staining for F480 (pan-macrophage marker), and either Cd40 (pro-inflammatory activation marker) or Ym1 (tolerogenic macrophage marker). Nerves with stained with pan-neuronal β-III Tubulin, with subtype specific staining using ChAT (choline acetyltransferase) and neuronal nitric oxide synthase (Nos1). Enteric neuronal progenitor cells were visualized with Ngfr-p75 staining, and smooth muscle structures with α-smooth muscle action (α-SMA). XY maximum intensity projections were obtained using the Olympus FluoView software, following consistent acquisition settings across all experimental conditions (control, PB, TNF-α). Quantitative fluorometry was performing using the corrected total fluorescence (CTF) method in FIJI/Image J using previously established protocols^19, 20, 45^. For each image, integrated density, area, and mean background fluorescence were measured for each ROI. CTF was calculated using the following formula:

CTF = Integrated Density – (Area of ROI × Mean Background Fluorescence)

To account for variations in cell number, CTF was divided by the number of nuclei (DAPI-positive cells) in each image, yielding an average CTF per cell. These CTF values were then normalized to the mean value from the control group to allow for relative comparison across conditions. Each experimental condition was replicated across multiple biological samples (n ≥ 5).

To assess macrophage activation, iBMDMs were co-stained for F480 (pan-macrophage marker) and Cd40 (pro-inflammatory activation marker) and imaged using standardized acquisition settings across all experimental groups. Quantification was performed using the same CTF method in ImageJ. For each image, Cd40 CTF was divided by the corresponding F480 CTF to generate a normalized Cd40/F480 fluorescence ratio, representing the degree of pro-inflammatory polarization relative to total macrophage content. Similar quantification was performed for Ym1/F480, to identify tolerogenic macrophage populations. Neuronal density changes with inflammatory insult were quantified via CTF of β-III Tubulin, ChAT, Nos1 and p75, while smooth muscle architectural changes were quantified via CTF of α-SMA. Each experimental condition was replicated across multiple biological samples (n ≥ 5).

### 2.6 Evaluation of contractility in bioengineered colon assembloids

Physiological contractility of bioengineered colon assembloids were assessed using a custom-built organ bath system^19, 20^. Each assembloid was mounted by securing one luminal end to a fixed reference post and the other to the arm of a force transducer (FT20, Harvard Apparatus). The organ bath was filled with DMEM buffered with 1 M HEPES and maintained at 37°C. A 30% passive stretch was applied using a micromanipulator and maintained throughout the experiment to record isometric force. Assembloids were equilibrated under these conditions for 40–60 minutes. Baseline force post-equilibration was arbitrarily set to zero to allow for direct comparison of pharmacologically induced changes in contractility.

To assess neural responsiveness and muscle function, assembloids were treated with the following agents: Acetylcholine (ACh, 1 μM/mL) to induce cholinergic contraction; Tetrodotoxin (TTX, 1 μM/mL) to reversibly inhibit neuronal sodium channels and block neuronal activity.

Pharmacologic agents were added directly to the bath, and contractile responses were recorded. Force traces were analyzed for peak amplitude and area under the curve (AUC) to quantify physiological responses. Each experimental condition was replicated across multiple biological samples (n ≥ 5).

### 2.7 Evaluation of macrophage mono-culture exposure to inflammatory stimuli

To initiate inflammatory insult on macrophage mono-cultures, iBMDMs were seeded and allowed to adhere overnight in densities ranging from 3,000 – 8,000 cells/mL. Following adhesion, on Day 1, macrophages were exposed to 100µM PB or 100ng/mL TNF-α for 24 hours. iBMDMs were assessed at different time points: ‘Acute’ was collected immediately after 24 hours of exposure (Day 2). To assess recovery following one-time, 24hr exposure to PB or TNF-α, removal of insult was followed by recovery in freshly supplemented growth media. Recovery was assessed longitudinally at two time points: 3 days post removal, or 5 days post removal of PB or TNF-α. Both these recovery time points were given the nomenclature of “Day 3 Chronic” to mean 3 days of recovery to assess lingering effects of inflammation, or “Day 5 Chronic” to mean 5 days of recovery to assess lingering effects of inflammation.

Macrophage activation in response to PB or TNF-α treatment was evaluated in multiple ways: (1) Gene expression analysis following methods outlined in Section 2.5.1, comparing time-matched untreated macrophages to PB or TNF-α treated macrophages. (2) Immunofluorescent co-staining of macrophages with pan-macrophage marker F480 and inflammatory marker Cd40, with CTF based quantitative fluorometry; (3) Cytokine profiling of cell culture supernatants following methods outlined in Section 2.5.3.

Media was also collected from iBMDMs following acute exposure to PB (Day 2), sterile filtered and termed as ‘PB-conditioned media (PB-CM)’. This media was used in future experiments, to mimic an inflammatory insult to cultured enteric neuronal progenitor cells.

### 2.8 Effect of inflammatory insult and prolonged macrophage-derived factors on enteric neuronal progenitor cell (IM-FEN) proliferation and identity

IM-FENs (8000 cells/mL) were allowed to adhere onto fibronectin-coated dishes overnight at 33°C, prior to initiation of differentiation. Once cells were adhered, progenitor IM-FENs were maintained at 33°C with no exposure (control) or constant inflammatory macrophage-derived conditioned media (PB-CM) for 7 days. Cells were fixed, and immunostained with the proliferation marker, Ki67 following immunofluorescence protocols.

The primary antibody, Ki67 (Alexa Fluor 488) was used to mark active cell proliferation. Ki67^+^ cells were identified by the presence of green fluorescence within DAPI (blue fluorescence) counterstained nuclei and were quantified manually using previously demonstrated protocols^46^. Analysis was performed to determine the number of actively proliferating cells (Ki67 antigen in the nuclei) versus the total cell number (number of nuclei) to determine the percentage of proliferating cells. Exposed cells were compared to controls. Each experimental condition was replicated across multiple biological samples (n ≥ 5).

IM-FENs were characterized using flow cytometry for Ngfr-p75 expression^47^, by adapting previously used flow analysis protocols^19, 20^. Briefly, IM-FENs were collected and resuspended in a flow cytometry amenable buffer (PBS + 2% FCS) at a concentration of 200,000 cells/ml. Cells were incubated with a p75-AF647 antibody, with a matched isotype control. Cells were then processed through a flow cytometer (Attune NxT, Waltham, MA, USA). Flow cytometry analysis was completed with FlowJo (Ashland, OR). Polygon gates were drawn on isotype flow cytometry graphs to establish a background cut off gate at 0.5% (gating strategy is presented in **Supplementary Figure 6**). The same gate was applied to antibody-stained cells to establish a percent positive population. Each experimental condition was replicated across multiple biological samples (n ≥ 3).

### 2.9 Effect of inflammatory insult and prolonged macrophage-driven factors on IM-FEN differentiation

IM-FENs (8000 cells/mL) were allowed to adhere onto fibronectin-coated dishes overnight at 33°C, prior to initiation of differentiation. Once cells were adhered, enteric neuronal progenitor cells were either maintained as untreated controls, or exposed to PB-CM in its progenitor state prior to differentiation for 24hrs, initiating the inflammatory insult in the progenitor cell state. Differentiation was initiated by increasing the temperature to 39°C on Day 2, supplemented with neural differentiation media. Cells were differentiated as untreated controls, or under the constant prolonged exposure of PB-CM over 5 additional days up to Day 7. Ability or changes in differentiation with prolonged PB-CM exposure was characterized using immunostaining for differentiated neuronal markers including β-III Tubulin, ChAT, and Nos1 and CTF-based quantification. Comparisons were drawn between untreated controls and PB-CM treated IM-FENs. Similar quantitative analyses were also carried out using gene expression analysis, to assess changes in PB-CM treated IM-FENs.

## 3. RESULTS AND DISCUSSION

### 3.1 Formation of a bioengineered 3D colon assembloid incorporating neural, smooth muscle, and immune lineages

Colonic motility arises from the muscularis externa of the colon, the outer layers that comprise smooth muscle cells, an integrated neuronal network and tissue-resident macrophages. To study functional consequences of enteric neuro-inflammation on colonic motility, we bioengineered the colonic muscularis externa. The bioengineered assembloid combined three key cell types within two layers of collagen hydrogel: (i) immortalized colonic smooth muscle cells (icSMCs) as the smooth muscle component that generates the forces relevant to colonic motility; (ii) immortalized enteric neuronal progenitor cells (IM-FENs) that differentiate within the bioengineered construct to form mature neuronal networks; and (iii) immortalized macrophages (iBMDMs) to add immune competency. The base inner layer contained all three cell types, and was cast loosely around a luminal post to define the hollow lumen. A second outer layer contained smooth muscle cells only, and enveloped the base layer and allowed to gel over 30 minutes. **Figure 1A** outlines the sequential cell seeding process into the collagen hydrogel, and subsequent self-assembly into concentric layered structures. Bioengineered colon assembloids matured over 1-7 days, enabling integrated neuro-immune structures and concentrically aligned smooth muscle architecture (**Figure 1B**). Compaction of the collagen hydrogel also occurred during the self-assembly process (>75% compaction over 3 days; **Figure 1C**).

**Figure 1.**
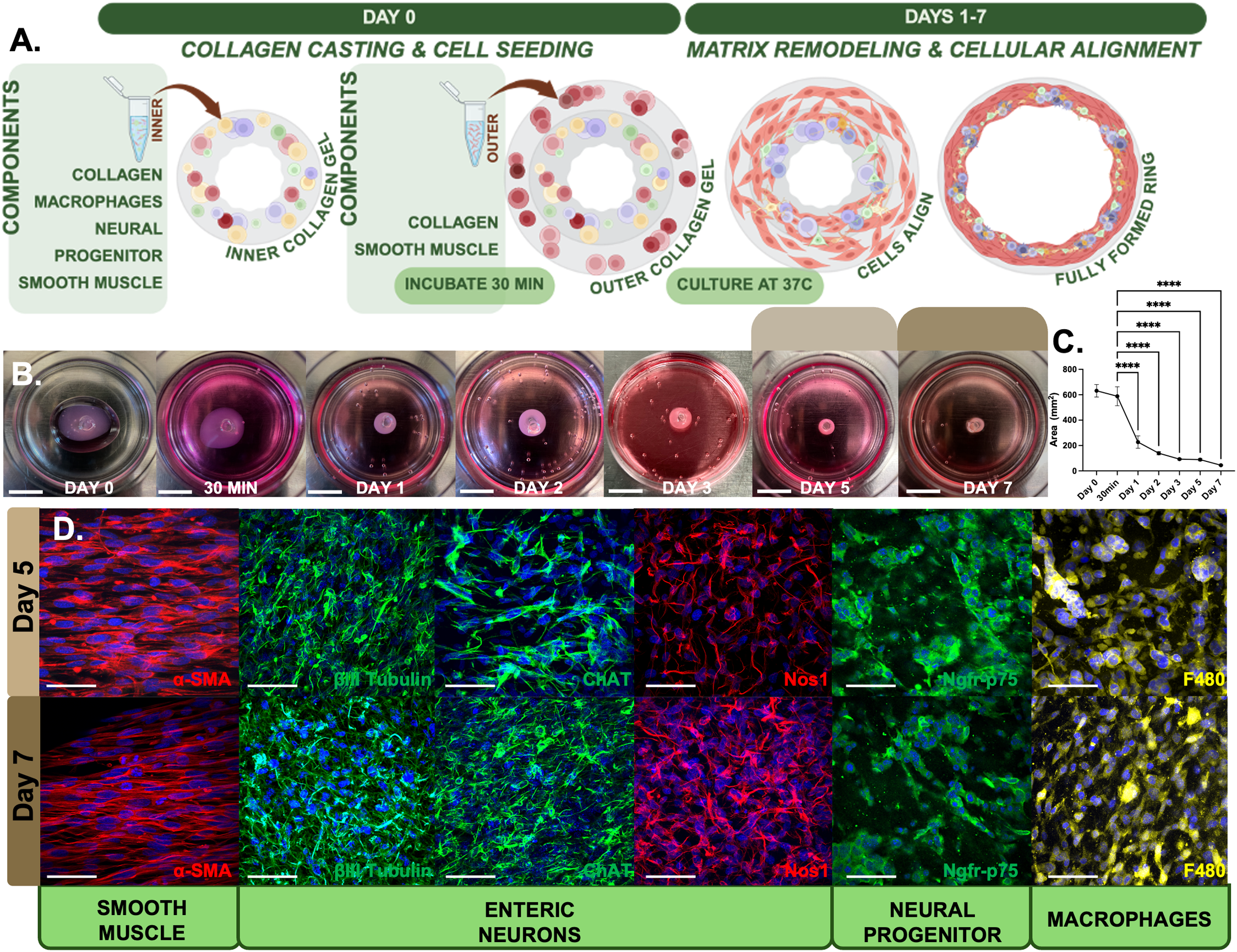
Multicellular 3D bioengineered colon assembloid integrating smooth muscle, neural, and macrophage immune cells. A) Schematic of the sequential collagen casting and cell seeding procedure. Layer 1 contained immortalized murine bone marrow-derived macrophages (iBMDM), immortalized murine fetal enteric neural progenitor cells (IM-FEN), and immortalized murine colonic smooth muscle cells (icSMC). These cells were embedded in an inner collagen gel, cast around a silicone post placed to define a hollow lumen. Once the first gel layer formed (∼15minutes), a second collagen layer was cast, laden with icSMCs alone (outer collagen gel). Constructs were cultured for 7 days at 37°C to self-assemble into a ring-like structure with a defined hollow lumen. B) Representative photographs of bioengineered colon assembloids during directed layer-by-layer assembly, and every day from days 1-7. A central post defining the hollow luminal space is still visualized in these photographs, with the bioengineered circular colon assembloid around it. Scale bar = 15 mm (C) Quantification of bioengineered colon assembloid area over time with progressive compaction. ****p<0.0001, ordinary one-way ANOVA, n>14. (D) Wholemount immunofluorescence staining at days 5 and 7 demonstrating cellular markers: α-SMA (α-smooth muscle actin), βIII-tubulin (neuronal tubulin), ChAT (choline acetyltransferase), and Nos1 (neuronal nitric oxide synthase), Ngfr-p75 (receptor for the neural growth factor p75 labeling neural progenitor cells), F4/80 (pan-macrophage marker). Scale bar = 20 µm.

Spatial and cellular maturation and organization within the bioengineered colon assembloids was confirmed via whole mount immunofluorescence (**Figure 1D**). By day 5, α-SMA^+^ smooth muscle cells were concentrically aligned, similar to the structural organization of smooth muscle cells in the colon^48^. Differentiated neurons derived from neuronal progenitor cells robustly expressed βIII-tubulin^+^, organized in reticular network-like structures similar to their organization within the myenteric plexus of the colon^49, 50^. Differentiated neuronal subtypes were further identified via stains specific to the synthesis of neurotransmitters, namely choline acetyltransferase (ChAT) and nitric oxide synthase (Nos1)^51^. Interestingly, our stains also demonstrated that bioengineered colon assembloids retained a population of neuronal progenitor cells in their progenitor state, evidenced by Ngfr-p75^+^ staining, similar to tissue resident adult enteric neuronal stem cells present in the human and mouse colons^52, 53^. Macrophage presence was additionally verified by F4/80^+^ expression interspersed within the bioengineered structure.

### 3.2 Bioengineered colon assembloids have distinct gene expression changes to stimulated neuro-inflammation

We next assessed the transcriptional consequences of inflammatory exposure in the bioengineered colon assembloids. We stimulated enteric neuro-inflammation using two different methods commonly associated with functional gastrointestinal disorders. The first stimulant was Pyridostigmine bromide (PB), a toxic exposure associated with Gulf War Illness^19, 20^, resulting in severe colonic motility disruption. The second stimulant we used in this study was Tumor Necrosis Factor-alpha (TNF-α), central to enteric neuro-inflammation in inflammatory bowel diseases like Crohn’s Disease^21^, where colonic motility is similarly disrupted. **Figure 2A** displays a timeline for when inflammatory stimuli are added to bioengineered colon assembloids on Day 3 once their compaction and self-assembly is complete. PB or TNF-α stimulation occurred for 24hrs (Day 3-4), following which an immediate response to inflammation was assessed (acute; Day 4). Once inflammatory stimulants were removed, bioengineered colon assembloids were allowed to recover for 3 additional days, and any chronic lingering effects of inflammation was assessed (chronic; Day 7). The recovery time window of 3 days was chosen based on previous studies assessing the dynamics of macrophage activation^43, 44^. All comparisons were made to untreated control bioengineered colon assembloids that were time-matched in culture.

**Figure 2.**
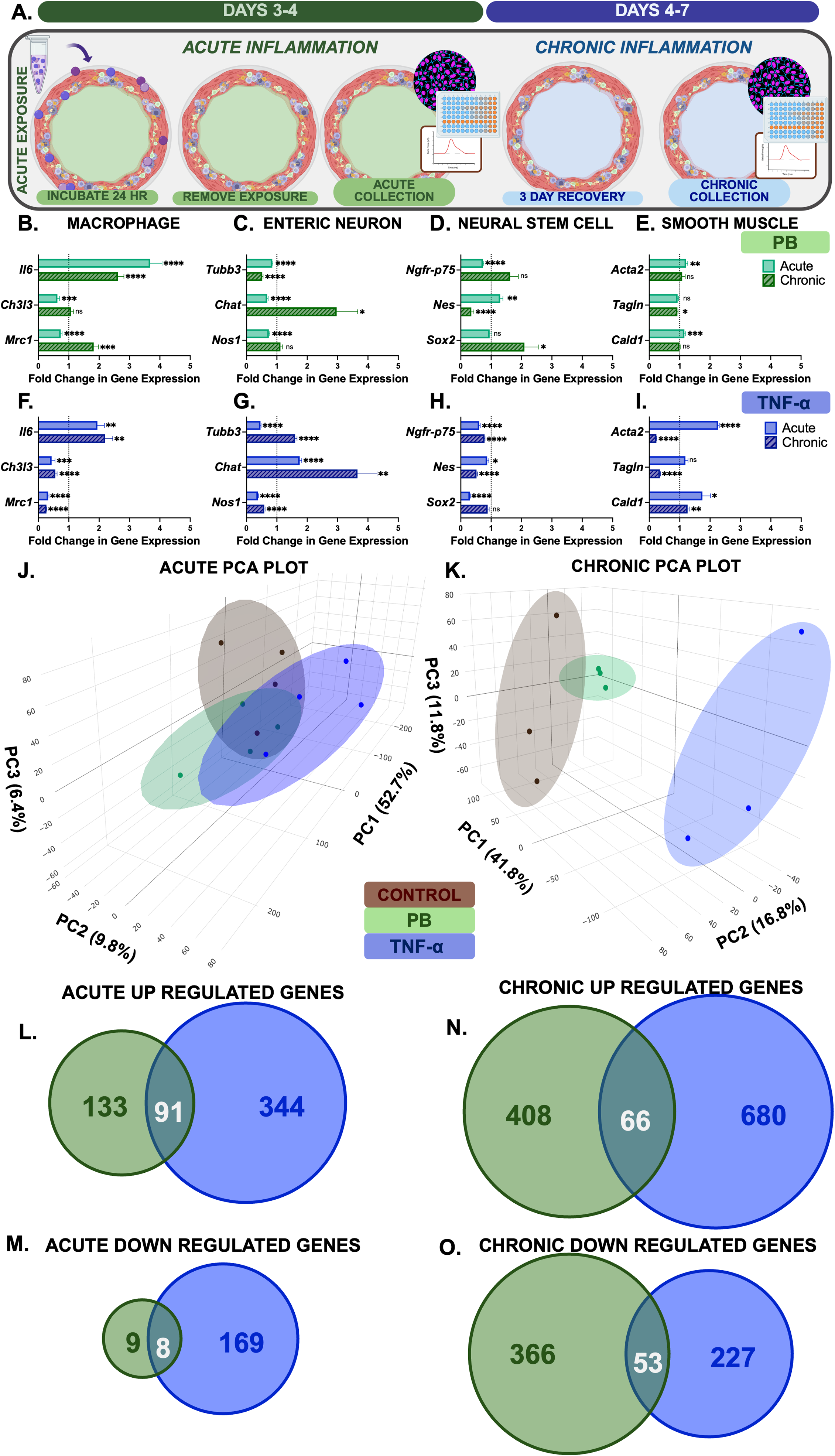
Transcriptomic changes in bioengineered colon assembloids induced by exposure to Pyridostigmine bromide (PB) or Tumor necrosis factor (TNF-α). A) Experimental timeline demonstrating timeline of inflammatory insult initiation at Day 3, and removal at Day 4; Acute effects were evaluated on Day 4; Chronic lingering effects of inflammation were evaluated after 3 days of recovery, on Day 7. (B–E) Gene expression analysis in PB-exposed bioengineered colon assembloids for macrophage (*Il6, Ch3l3, Mrc1*) (B), enteric neuron (*Tubb3, Chat, Nos1*) (C), neural stem cells (*Ngfr-p75, Nes, Sox2*) (D), and smooth muscle (*Acta2, Tagln, Cald1*) (E) markers, comparing acute and chronic conditions to untreated control. (F–I) Corresponding gene expression changes in TNF-α exposed rings for macrophage (F), enteric neuron (G), neural stem cell (H), and smooth muscle (I) markers. The dotted line at 1 represents the normalized level of gene expression for time-matched control colon assembloids. Data are presented as mean ± SEM. ****p<0.0001, ***p<0.001, **p<0.01, *p<0.05, ‘ns’ = not significant, one sample t-test, n>3. (J, K) Three-dimensional principal component analysis (PCA) of bulk RNA-seq showing clustering of control (brown), PB-treated (green), and TNF-α treated (blue) bioengineered colon assembloids under acute (n=4; J) and chronic (n=3; K) conditions. Each dot represents an individual biological replicate. % values indicate the proportion of total variance within each principal component. (L, M) Venn diagram comparing overlapping of significantly upregulated (L) and downregulated (M) genes between acute PB and TNF-α exposure. (N, O) Venn diagram comparing overlapping of significantly upregulated (N) and downregulated (O) genes in lingering chronic PB and TNF-α induced inflammation.

#### 3.2.1 PB-induced gene expression changes in bioengineered colon assembloids mimic key aspects of those observed in Gulf War Illness mouse colons

Acute PB exposure significantly increased the gene expression of *Il6* (**3.68-fold**), with a subsequent drop to 2.62-fold chronically, indicating mild but incomplete recovery (**Figure 2B**). The elevated low-grade inflammation in bioengineered colon assembloids was in line with observations from mouse models of persistent enteric neuro-inflammation in Gulf War Illness^20^. Markers of reparative macrophages, *Mrc1* and *Ch3l3*, were acutely downregulated, with a partial restoration following recovery.

Acute PB exposure also resulted in a reduction of neuronal gene expression compared to untreated bioengineered colon assembloids (**Figure 2C**). Specifically, *Tubb3* (**0.85-fold; ****p<0.0001**), *Chat* (**0.68-fold; ****p<0.0001**), and *Nos1* (**0.73-fold; ****p<0.0001**) were all significantly decreased, indicating impaired neuronal programming. Interestingly, upon recovery, *Chat* expression trended towards significant upregulation (**2.97-fold; \*\*\*\**p*<0.0001**). These findings in bioengineered colon assembloids also mimicked studies in Gulf War Illness mouse models, where one-time PB exposure chronically destroyed enteric neuronal integrity and altered neuronal subtype balance^20^. Neuronal recovery, despite subtype imbalance, was supported by increased gene expression of *Ngfr-p75* and *Sox2* (**Figure 2D**), in line with adult enteric neo-neurogenesis following inflammatory insult^54, 55^, captured within bioengineered colon assembloids. Interestingly, PB exposure did not alter the gene expression of smooth muscle markers (**Figure 2E)**, also in line with our previous observations in mouse models^20^.

#### 3.2.2 TNF-α–induced gene expression changes are distinctly different compared to PB-induced changes in bioengineered colon assembloids

Bioengineered colon assembloids exposed acutely to TNF-α exhibited strong pro-inflammatory activation (**Figure 2F**), with significant upregulation of *Il6* (1.94-fold; **\*\*p<0.01**). Reparative macrophage markers, *Mrc1* and *Ch3l3,* were reduced (**0.32-** and **0.44-fold**, **\*\*\*p<0.001**) compared to control, with no evident recovery in gene expression levels. This is unsurprising, because TNF-α is a potent inflammatory mediator in macrophages, including in inflammatory bowel disease^21^.

Neuronal damage and mild recovery were observed in bioengineered rings with TNF-α treatment, with acute downregulation of *Tubb3* and *Nos1,* but not *Chat* (**Figure 2G**). Markers of neuronal stemness were more broadly suppressed with TNF-α treatment, compared to PB, both acutely and chronically despite recovery. *Sox2* expression partially recovered to 0.89-fold (**\*\*\*\*p<0.0001)** but remained significantly lower than control (**\*p<0.05; Figure 2H**). *Ngfr-p75* slightly increased to 0.78-fold (**\*\*p<0.01**), while *Nes* declined further to 0.52-fold (**\*\*\*\*p<0.0001**) compared to acute rings, demonstrating complex and marker-specific neural stem cell regulation over chronic TNF-α induced inflammation.

Acute TNF-α exposure induced pronounced upregulation of smooth muscle contractile genes *Acta2* (**2.27-fold; ****p<0.0001**), *Cald1* (**1.75-fold; *p<0.05**), and *Tagln* (**1.19-fold; ns),** compared to control rings (**Figure 2I**). In line with observations from experimental mouse models of inflammatory bowel disease, TNF-α exposure significantly alters expression of thin filament contractile elements within smooth muscle like Caldesmon (*Cald1*) and Smoothelin (*Tagln),* that ultimately result in impaired colonic contractility and motility^56, 57^. Bioengineered colon assembloids capture this important nuance, where TNF-α treatment resulted in changes in the smooth muscle compartment within the bioengineered colon, while PB did not.

### 3.3 Unbiased transcriptomics captures distinctions in PB and TNF-α induced enteric neuro-inflammation within bioengineered colon assembloids

In order to capture the differences in assembloid response to both acute and chronic PB or TNF-α stimulation, we performed three-dimensional principal component analysis (3D PCA) on bulk RNA-seq data from treated and untreated bioengineered colon assembloids. The acute 3D PCA plot (**Figure 2J**) illustrates distinct clustering of the three groups along principal components 1, 2, and 3, which collectively explain 68.9% of the total variance (PC1 = 52.7%, PC2 = 9.8%, PC3 = 6.4%). Samples from PB- and TNF-α treated groups separate clearly from controls, demonstrating transcriptomic alterations in response to inflammatory challenge within the bioengineered colon assembloids. These transcriptional distinctions persisted into the chronic phase, as shown in the chronic PCA plot (**Figure 2K**), where the three groups again clustered separately, with total variance explained reaching 70.4% (PC1 = 41.8%, PC2 = 16.8%, PC3 = 11.8%). This continued separation suggests lasting and distinct transcriptional remodeling despite removal of the initial stimulus.

The upregulated gene sets (**Figure 2L**) revealed both overlapping and distinct genes: a core subset of 91 genes was commonly upregulated by both PB and TNF-α, likely indicative of shared inflammatory pathways. Additionally, TNF-α uniquely upregulated 344 genes, while PB uniquely upregulated 133 genes, reflecting the operation of stimulus-specific transcriptional programs. Similarly, the downregulated gene sets (**Figure 2M**) revealed an overlap of 8 genes, with TNF-α driving broader suppression of 169 unique genes compared to PB’s 9 unique genes. After recovery, chronic lingering effects of inflammation led to even more divergent transcriptional responses. A total of 408 genes were uniquely upregulated by PB, and 680 by TNF-α, with 66 genes commonly upregulated between the two treatments (**Figure 2N**). Similarly, 366 genes were uniquely downregulated by PB, 227 by TNF-α, and 53 genes were shared (**Figure 2O**).

### 3.4 Gene Ontology and biological process analyses reveal shared and distinct mechanisms of PB and TNF-α induced inflammation and impaired neurogenesis

Noting that PB and TNF-α exposure generated some distinct inflammatory signatures within bioengineered colon assembloids (differentially regulated genes also visualized as volcano plots in **Supplementary Figure 1A, B**), we first explored unbiased RNA-Seq analysis to further determine distinctions. **Supplementary Figure 1** also demonstrates distinct genes that drive unique but strong immunologic signatures upon PB and TNF-α exposure, with distinctions between the two exposures broken down further in **Supplementary Figure 2**. The involvement of shared and distinct gene signatures was further highlighted via unbiased gene ontology and biological process (GO; BP) enrichment analyses (**Figure 3A-D**). Both stimuli (PB and TNF-α) resulted in positive regulation and activation of innate immune response within the bioengineered colon assembloids. Within the colonic muscularis externa, macrophages are key regulators of immune response to inflammation and stress, resulting in changes in colonic motility and neuronal health^16^. Transcriptomic changes confirmed that macrophages included within the assembloids were able to respond to inflammatory stimuli like PB or TNF-α, the functional consequences of which we explore later in the study.

**Figure 3.**
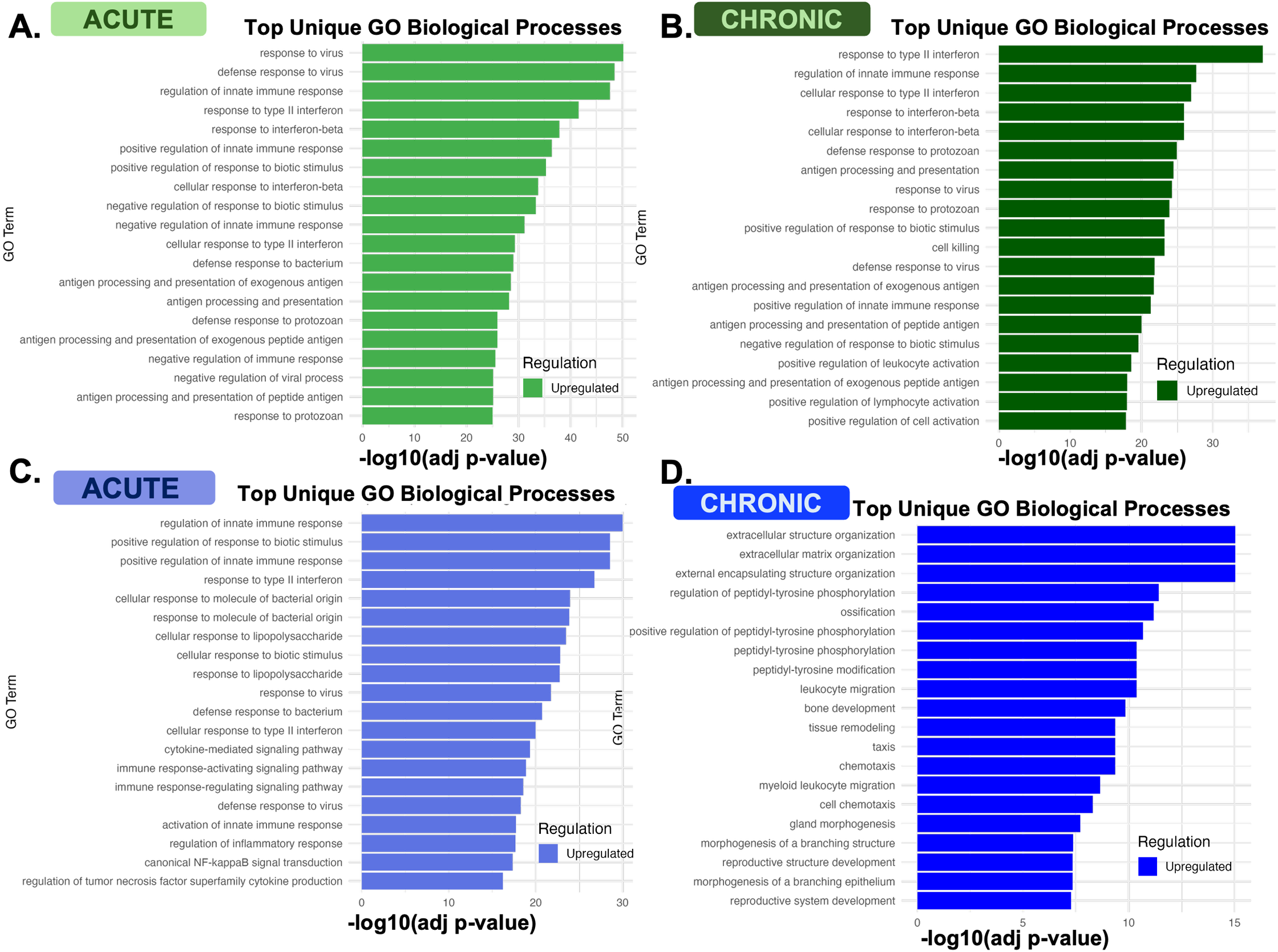
Transcriptomic profiling and biological process enrichment following exposure to PB and TNF-α in bioengineered colon assembloids. (A, B) Top unbiased Gene Ontology (GO) Biological Processes (BP) significantly enriched in PB-treated assembloids acutely (A) and chronic effects of one-time PB treatment (B). (C, D) Top unbiased GO BP significantly enriched in assembloids treated with TNF-α and assessed acutely (C) and for chronic effects (D). Bars represent the –log10 adjusted p-value for each GO term.

Despite the seeming similarities of inflammation responses initiated by PB and TNF-α, a deeper investigation of enriched gene sets highlighted subtle differences in molecular mechanisms underlying the exposures (**Figure 3A-D, Supplementary Figure 3**). For each identified pathway, a z-score was calculated based on the proportion of significant genes within the pathway.

The toxic exposure-based stimulant PB resulted in pronounced interferon-gamma based inflammatory responses in both acute and chronic settings (**Figure 3A-B; Supplementary Figure 3A,B)**, in line with many observations in Gulf War veterans that have higher serum levels of interferon-gamma^58, 59^. In fact, interferon-gamma based neuro-inflammation is documented to drive many neurological disorders central to Gulf War Illness, priming the innate immune system to sustain neuro-inflammation chronically^60^.

In addition to inflammation via interferon-gamma based response, acute TNF-α exposure also drove inflammation via NF-κB signaling, with stronger enrichment of cellular response to interleukin-1 (**Figure 3C-D**; **Supplementary Figure 2D, Supplementary Figure 3C**). TNF-α and interleukin-1 are elevated in functional gastrointestinal disorders and inflammatory bowel disease^61,62^. Interestingly, chronic TNF-α exposure strongly enriched chemotaxis (**Supplementary Figure 3D**), where persistent immune cell recruitment contributes to tissue remodeling and fibrosis^63^. These findings were also supported by changes in regulatory transcription factor-gene interaction maps (**Supplementary Figure 4**) between the two inflammatory stimuli.

Overall, unbiased transcriptomic analysis revealed that bioengineered colon assembloids were capable of responding to inflammatory stimuli, initiated either by PB exposure or by TNF-α exposure. Importantly, even after removal of the initial insult (chronic conditions), each exposure sustained slightly divergent signatures during recovery: PB-driven responses remained centered on interferon-gamma mediated neuroinflammation and impaired neurogenesis, while TNF-α shifted toward extracellular matrix remodeling and fibrotic pathways, mimicking key aspects of published work in GWI and IBD.

### 3.5 Cytokine profiling reveals distinct protein levels changes in bioengineered colon assembloids under neuro-inflammatory stimuli

Secreted cytokines are key orchestrators of tissue damage downstream of inflammatory initiation. Given the prevalent, yet subtly distinct, inflammatory gene signatures in bioengineered colon assembloids with PB and TNF-α exposure, we next investigated changes in a panel of secreted cytokines that amplify inflammation^64–68^, mediate neural repair^69, 70^ or are associated with tissue damage within the colonic muscularis externa^71^. PB and TNF-α treatment demonstrated partially distinct profiles (**Figure 4**).

**Figure 4.**
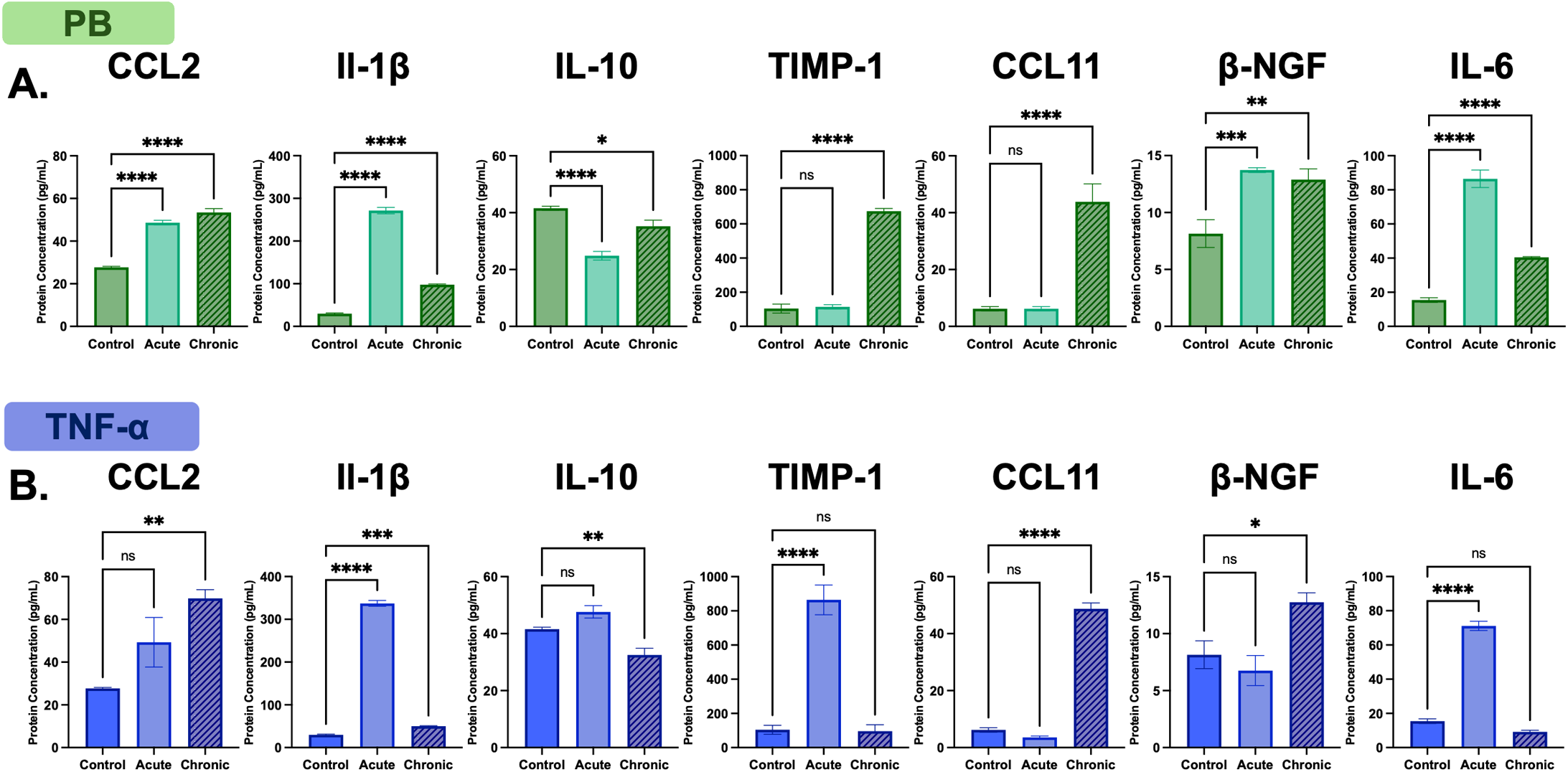
Cytokine profiling of bioengineered colon assembloids following exposure to PB and TNF-α. (A, B) Multiplex Luminex quantification of inflammatory and regulatory cytokines from bioengineered colon assembloids under untreated control, or PB or TNF- α treatment for 24hrs. Treatment was assessed either acutely (immediately upon 24hr exposure), or lingering chronic effects were assessed following a 3-day recovery after initial inflammatory insult. Analyzed cytokines include: CCL2 (monocyte chemoattractant protein-1), IL-1β (interleukin-1 beta), IL-10 (interleukin-10), TIMP-1 (tissue inhibitor of metalloproteinases-1), CCL11 (eotaxin), β-NGF (beta-nerve growth factor), and IL-6 (interleukin-6). A) Cytokine levels following PB treatment in bioengineered colon assembloids. B) Cytokine levels following TNF-α treatment in bioengineered colon asembloids. Data are presented as mean ± SEM. ****p<0.0001, ***p<0.001, **p<0.01, *p<0.05, ‘ns’ = not significant; ordinary one-way ANOVA; n>3.

#### 3.5.1 Single PB exposure results in chronic low-grade neuro-inflammation and pathological tissue remodeling

Acute exposure to PB triggered a robust pro-inflammatory response in bioengineered colon assembloids, characterized by elevated CCL2, IL-1β, and IL-6 (**\*\*\*p<0.001; Figure 4A; ‘Acute’**) and decreased IL-10 (**\*\*\*\*p<0.0001**). Cytokine production was also in-line with identified GO processes identified within bioengineered colon assembloids enriched following PB-exposure (**Supplementary Figure 3A**). This mirrors neuro-inflammatory profiles reported in the muscularis externa of GWI-affected colons^19, 20^.

Following removal of PB and subsequent recovery, the lingering inflammatory environment in bioengineered colons assembloids evolved (noted as ‘Chronic’ in **Figure 4**): TIMP-1 and CCL11 levels increased (**\*\*\*\*p<0.0001**), reflecting pathological tissue remodeling commonly seen within the inflamed colon^72, 73^. Low-grade, unresolved inflammation persisted through elevated IL-6 and IL-1β, along with low IL-10 levels, similar to GWI^74^. Additionally, persistent elevation of β-NGF (**\*\*p<0.01**) was indicative of neuroinflammation driven altered neuronal plasticity^75^. Together, these findings showed that PB exposure induced a chronic low-grade inflammatory state in the macrophage-immune competent bioengineered colon assembloids.

#### 3.5.2 TNF-α induced secretome partially reflects inflammation resolution with time

Acute TNF-α exposure prompted pro-inflammatory cytokine secretion (IL-1β, IL-6) in bioengineered colon assembloids (**Figure 4B; ‘Acute’**). Interestingly, TNF-α (but not PB) immediately increased TIMP-1, a sign of acute pathological tissue remodeling. Following the removal of TNF-α, bioengineered colon assembloids progressed partially towards recovery (**Figure 4B; ‘Chronic’**), with plummeting levels of IL-1β, IL-6 and TIMP-1. Mild signatures of neuro-inflammation were still persistent, including a potential recovery, with elevated CCL11 and β-NGF.

Overall, bioengineered colon assembloids continued to capture key aspects of inflammatory cytokine landscapes and remodeling processes similar to that observed in the colon, that contribute to motility dysfunction within the muscularis externa^76, 77^.

### 3.6 Wholemount immunostaining reveals structural changes in bioengineered colon assembloids with neuro-inflammatory stimulation

Next, we used whole mount immunostaining to visualize structural changes that resulted from secreted factors and gene expression associated with neuro-inflammation, tissue remodeling, and neural plasticity. As the likely responder and orchestrater of inflammation, we first assayed macrophage numbers and phenotypes within the bioengineered colon assembloids. Overall neuronal network integrity was assessed using βIII-tubulin, with cholinergic subtypes visualized via ChAT (choline acetyltransferase), and inhibitory nitrergic subtypes using Nos1 (neuronal nitric oxide synthase). The continued presence of neuronal progenitor cells was also assayed, since these progenitor cells contribute to neuronal repair. Lastly, the effectors of motility, the smooth muscle cells were identified for their arrangement using α-smooth muscle actin to additionally note tissue structure integrity changes (**Figure 5**). Changes were compared between control untreated and PB or TNF-α stimulated bioengineered colon assembloids.

**Figure 5.**
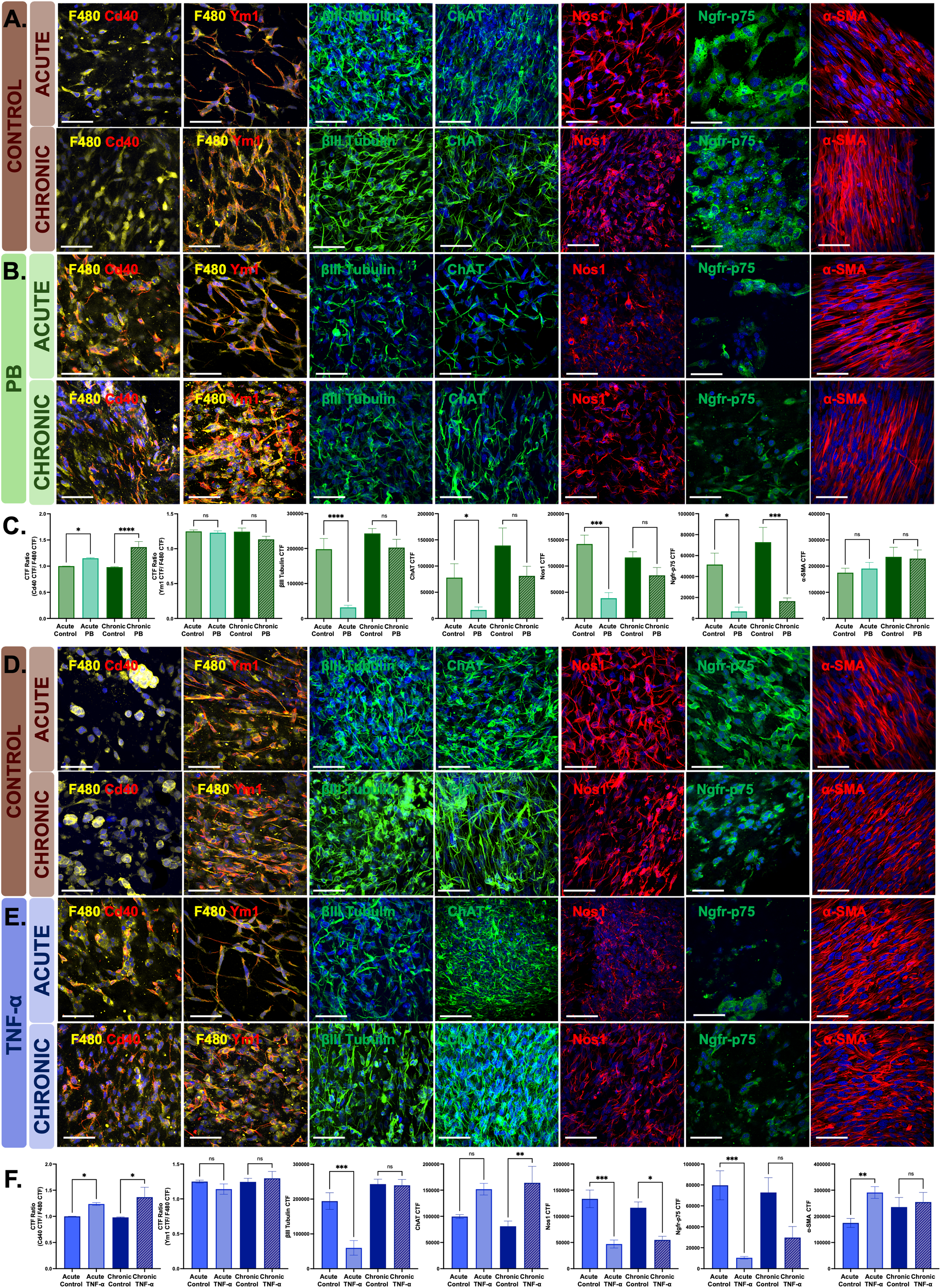
Wholemount immunostaining in bioengineered colon assembloids as untreated controls or following PB or TNF-α exposure. (A–C) Representative immunofluorescence images of control (A) and PB-exposed (B) bioengineered colon assembloids at acute (24hrs following stimulation) and chronic (3 days of recovery after initial inflammatory stimuli) timepoints. Scale bar = 20 µm. (C) Quantification of macrophage phenotypes (F4/80, yellow; Cd40, red; Ym1, red), neuronal subtypes (βIII-tubulin, ChAT, Nos1-all green), neural progenitors (Ngfr-p75, green), and smooth muscle (α-SMA, red) in control and PB-treated bioengineered colons under acute (24-hr exposure) and chronic (3-day recovery) conditions. (D–F) Representative immunofluorescence images of control untreated (D) and TNF-α (E) treated bioengineered colon assembloids at acute and chronic timepoints. Scale bar = 20 µm. (F) Quantification of control and TNF-α treated rings at acute and chronic time-points. Data are presented as mean ± SEM. ****p<0.0001, ***p<0.001, **p<0.01, *p<0.05, ‘ns’ = not significant; ordinary one-way ANOVA; n>5.

#### 3.6.1 PB exposure drives sustained inflammation with impaired neural regeneration

Inflammatory Cd40^+^ macrophages significantly increased with PB-treatment, rising 15% acutely and staying elevated by 37% in chronic conditions compared to time-matched untreated controls (**\*p<0.05; Figure 5A-C**). No changes were observed in reparative Ym1^+^ macrophages, consistent with its associated gene expression of *Ch3l3* (**Figure 2B**) and relatively stable levels of IL-10 (**Figure 4A**).

Acute and sustained inflammation lead to neuronal degeneration^19, 78, 79^. Structural damage of the neuronal network was evident with visual changes in βIII-tubulin organization (84% reduction with PB, compared to untreated controls; **Figure 5A-C**). Cholinergic and nitrergic neuronal subtypes were also reduced (79% and 73% respectively; **\*p<0.05; Figure 5B, C**). Acute cholinergic nerve loss is unsurprising following PB treatment, since PB reversibly inhibits acetylcholine esterase, thereby directly targeting cholinergic nerves^80^. Sustained loss of neuronal integrity despite recovery (‘Chronic’ in **Figure 5B-C**) reflects a compromised neuronal repair system^81^. Consistent with this notion, PB treatment reduced neuronal progenitor cells by 87% (Ngfr-75; **\*p<0.05; Figure 5A-C**) with no recovery, contributing to the disarrayed neuronal network.

Smooth muscle alignment and structural integrity largely remained unchanged, with only minor fluctuations (9% increase acutely, 3% decrease chronically; **Figure 5A-C**), in line with gene expression data (**Figure 2E**). This stability suggested that smooth muscle structural integrity was preserved despite neuroimmune disruption.

#### 3.6.2 TNF-α exposure drives distinct neuronal changes along with reactive smooth muscle hypertrophy

Similar to PB-exposure, TNF-α also increased inflammatory Cd40^+^ macrophages in bioengineered colon assembloids (23% acutely and 37% chronically compared to untreated controls; **\*p<0.05; Figure 5D-F**). Unchanging reparative Ym1 dynamics continued to suggest an impaired resolution of inflammation in the timeframes the bioengineered colon assembloids were examined.

TNF-α exposure also contributed to acute disintegration of neural network integrity (69% loss in βIII-tubulin fluorescence, **\*p<0.05, Figure 5D-F**). While cholinergic ChAT^+^ neurons were not affected with TNF-α like PB, a severe loss of nitrergic Nos1^+^ neurons occurred. The loss in nitrergic nerves is in line with therapeutic evidence that anti-TNF monoclonal antibodies aid in robust nitrergic neuronal recovery in experimental IBD models^82^. Interestingly, following recovery from TNF-α exposure, neuronal network integrity and ChAT^+^ neurons were fully restored (**‘Chronic’ TNF-α, Figure 5E**). This mild neurotrophic effect of TNF-α on enteric neurons, especially in co-culture systems where smooth muscle cells are present has been observed previously^83^, and replicated within bioengineered colon assembloids.

As expected, α-SMA expression increased acutely by 67% (**\*\*p<0.01**), consistent with massive increases in secreted TIMP1, and reactive smooth muscle hypertrophy observed in IBD^84, 85^. This contrasts with the stable α-SMA levels seen following PB exposure. During recovery, α-SMA expression returned to baseline, indicating a resolution of the hypertrophic response over time.

### 3.7 Inflammatory exposures impair colonic motility in a 3D bioengineered colon model

Altered colonic motility is a hallmark of both GWI^86^ and IBD^87, 88^, contributing significantly to functional gastrointestinal disorders and reduced quality of life^89–91^. Coordinated contractile function is a complex interplay of homeostatic macrophage immune activation, structural integrity of smooth muscle cells, intact neuronal architecture and balanced neuronal subtypes. In response to inflammatory stimuli in bioengineered colon assembloids, we observed pervasive changes in inflammatory macrophage activation (Cd40 enrichment in Figure 5; cytokine secretome in Figure 4; genotype in Figures 2-3), the structure of smooth muscle arrangement (like those observed in Figure 5E), and neuronal damage and imbalance in neurochemical coding (seen as imbalanced ChAT/Nos neurons in Figure 5). We hypothesized that these changes will alter colonic motility in PB or TNF-α exposed colon assembloids compared to control, untreated colon assembloids. To this end, we assessed contractile behavior in bioengineered colon assembloids using a force transducer-based organ bath system^19, 20, 92^. Bioengineered colon assembloids were stabilized in organ baths, and contractions were recorded following exogenous stimuli designed to interrogate both neuronal and smooth muscle contributions. Representative contractility traces from control and inflamed assembloids (PB- and TNF-α exposed) are shown in **Figure 6**, with time on the X-axis and force amplitude relative to baseline on the Y-axis.

**Figure 6.**
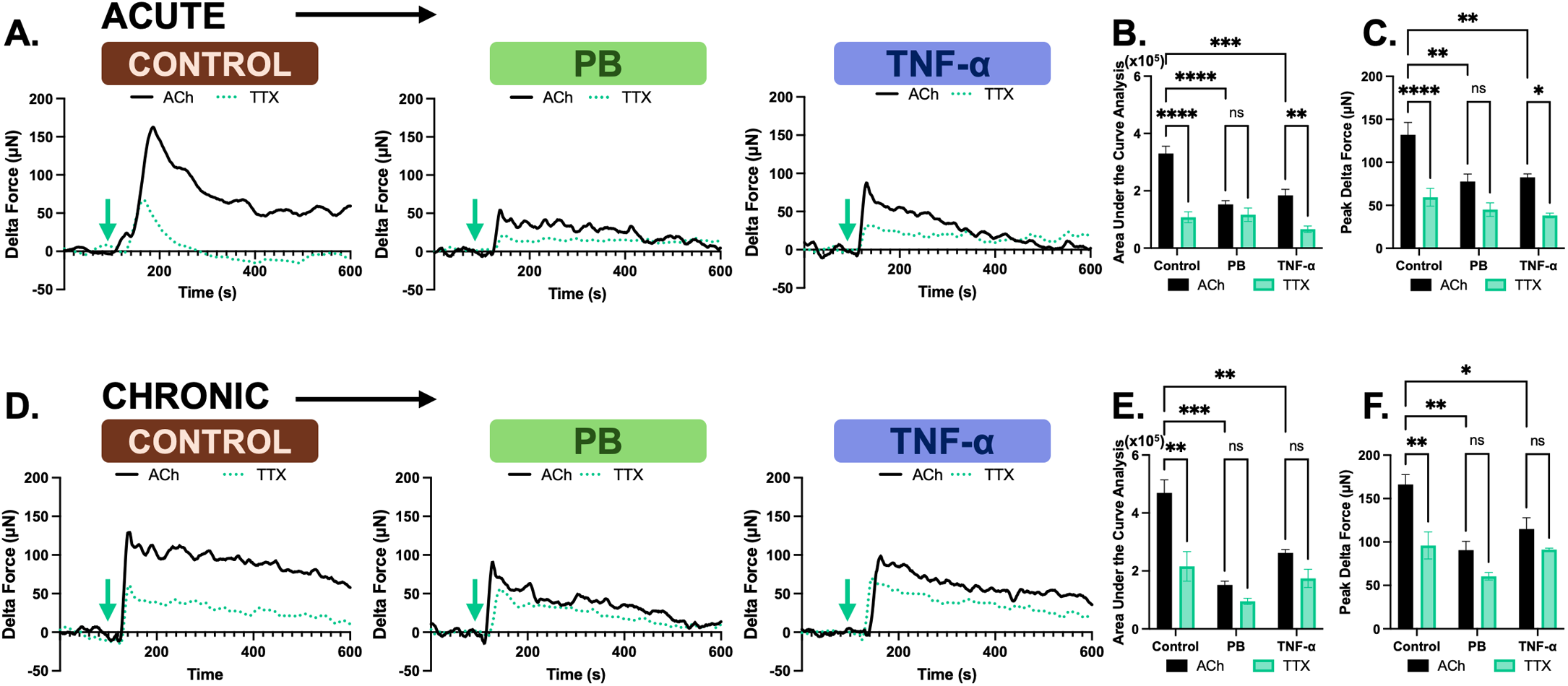
Inflammatory exposures impair neuronally mediated contractility in 3D bioengineered colon assembloids. Bioengineered colon assembloids were mounted in a force transducer-fitted organ bath, and contractile force was measured in response to exogenous addition of acetylcholine (ACh). ACh addition is indicated by the blue arrow, after the establishment of a stable baseline of force (arbitrarily set to zero). (A) Representative traces in response to 1 μM ACh (black traces) stimulation in control untreated or acute exposure to PB-, or TNF-α. Neuronal contributions were isolated by pre-treatment with 1 μM tetrodotoxin (TTX, blue dotted lines. (B) Quantification of contractile output as area under the curve for various treatment conditions under acute exposure. Data are presented as mean ± SEM. ****p<0.0001, ***p<0.001, **p<0.01, ‘ns’ = not significant; two-way ANOVA; n>5. (C) Quantification of contractile output as peak magnitude of force generated following ACh stimulation in acute conditions. Data are presented as mean ± SEM. ****p<0.0001, **p<0.01, *p<0.05, ‘ns’ = not significant; two-way ANOVA; n>5. (D) Representative traces in response to ACh stimulation in chronic bioengineered colons, maintained as untreated controls or 3 days recovered from PB or TNF-α treatment. (E) Quantification of contractile output as area under the curve for various treatment conditions under chronic exposure. Data are presented as mean ± SEM. ***p<0.001, **p<0.01, ‘ns’ = not significant; two-way ANOVA; n>3. (F) Quantification of contractile output as peak magnitude of force generated following ACh stimulation in chronic conditions. Data are presented as mean ± SEM. **p<0.01, *p<0.05, ‘ns’ = not significant; two-way ANOVA; n>3.

#### 3.7.1 Inflammation causes persistent impairment of colonic contractile function mediated by cholinergic neuronal activity

Contractile function was assessed by exogenous addition of acetylcholine (ACh; timing indicated by arrows in **Figure 6**). To isolate neuronal functionality in addition to the direct effect of ACh on smooth muscle cells, bioengineered colon assembloids were pre-treated with a nerve blocker, tetrodotoxin (TTX; green dotted traces in **Figure 6**). Control untreated bioengineered colon assembloids displayed robust ACh-induced contractions with an average peak force of 132.0 ± 0.8 µN (black trace, **Figure 6A, C**). Pretreatment with TTX significantly reduced contractile amplitude by 67% (green dotted trace, **\*\*\*\*p<0.0001; Figure 6B**), confirming the presence of a functional neuronal component capable of mediating contractions within bioengineered colon assembloids^92^.

Both PB and TNF-α treatment reduced ACh-induced contractile magnitude (53% lower in PB; 44% lower in TNF-α; **\*\*\*\*p<0.0001, ***p<0.001; Figure 6B, C**). Interestingly, acute structural loss of cholinergic nerves with PB treatment (but not TNF-α) manifested functionally as muted sensitivity to TTX pre-treatment (only 23% sensitive in PB-exposed assembloids, compared to 67% sensitivity in untreated controls and 64% sensitivity in TNF-α treated assembloids). Following removal of PB and recovery, contractile dysfunction persisted in chronic PB-treated assembloids (**Figure 6D-F**), similar to loss of contractility in GWI colons^19^. Only partial recovery of contractility was observed in chronic TNF-α treated assembloids (***p<0.001; **\*\*p<0.01**; **Figure 6E, F**), despite a recovery of structural neuronal integrity. The broader colonic dysmotility is further supported by immunostaining and gene expression data, where an imbalance in excitatory and inhibitory neuronal populations is evident, which can impair coordinated motility^20, 93, 94^. Especially in the TNF-α treated colons, a further contributing factor to diminished contractility is the reactive changes in smooth muscle that can alter muscle responsiveness, limiting contractile efficiency^95, 96^. Future testing will expand to different pharmacological agents capable of testing neuronal connectivity and muscarinic receptor sensitivity, for deeper characterization of the impact of neuro-inflammation on colonic contractility.

### 3.8 Inflammatory secretome from PB-exposed macrophages disrupts neural stem cell maintenance and impairs neuronal differentiation

While the 3D bioengineered colon assembloid model captured key aspects of the complex interactions between neuro-inflammation and motility, the presence of multiple interacting cell types and extracellular matrix components makes it difficult for finer mechanistic dissection. To disentangle the direct effects of inflammatory stimuli, such as PB and TNF-α, we first employed a simplified macrophage monoculture system. Macrophages treated with PB for 24hrs in cell culture released cytokines, creating a soluble inflammatory milieu supporting low-grade inflammation, notably with elevated IL-6 (**Supplementary Figure 6**). This inflammatory milieu was captured in the form of conditioned media derived from macrophages that were treated with PB for 24hrs (PB-CM). This resolution observed in macrophage monocultures contrasts with the persistent inflammation seen within 3D bioengineered colon assembloids, where complex multicellular interactions and extracellular matrix components likely sustain pro-inflammatory signaling and impede full recovery. Thus, while macrophages have an intrinsic capacity to resolve inflammation after PB exposure in vitro, the 3D microenvironment introduces additional regulatory factors that complicate resolution, highlighting the importance of studying inflammation in physiologically relevant contexts.

Next, we employed a monoculture model of immortalized p75^+^ enteric neuronal progenitor cells (IM-FENs)^39^, which possess the ability to differentiate into motor neuronal subtypes^92^. First, we directly exposed IM-FENs to PB for 24hrs to assess their intrinsic sensitivity to inflammatory insult (**Supplementary Figure 6**). While PB treatment induced some changes in proliferation and viability, the overall effects of PB on differentiation capability were modest. One-time exposure to PB in IM-FENs did not significantly alter the differentiation of IM-FENs into cholinergic or nitrergic neurons (**Supplementary Figure 6F-H**). These results implied that even though PB treatment induced a loss in the progenitor pool, differentiation capacity was only modestly impacted in monocultures.

However, our previous observations in mouse models of PB exposure^20^ and bioengineered colon assembloids (**Figure 5,6**) demonstrated significant damage in neuronal structural integrity resulting in functional dysmotility, implying that multi-cellular interactions may drive neuronal progenitor damage. Indeed, from PB mouse models, we observed inflammatory Cd40 macrophages in close proximity to p75+ enteric neuronal progenitor cells within the colonic myenteric plexus^20^. Therefore, we hypothesized that impairment in enteric neuronal progenitor differentiation might be macrophage mediated. To test this hypothesis, we next applied conditioned media collected from PB-treated macrophages (PB-CM; iBMDMs) to IM-FEN cultures (**Figure 7A**). Macrophages were only treated with PB for 24hrs, and media was collected acutely right away following exposure. IM-FENs were chronically exposed to PB-CM conditioned medium, to simulate ongoing low-grade enteric neuro-inflammation consistently.

**Figure 7.**
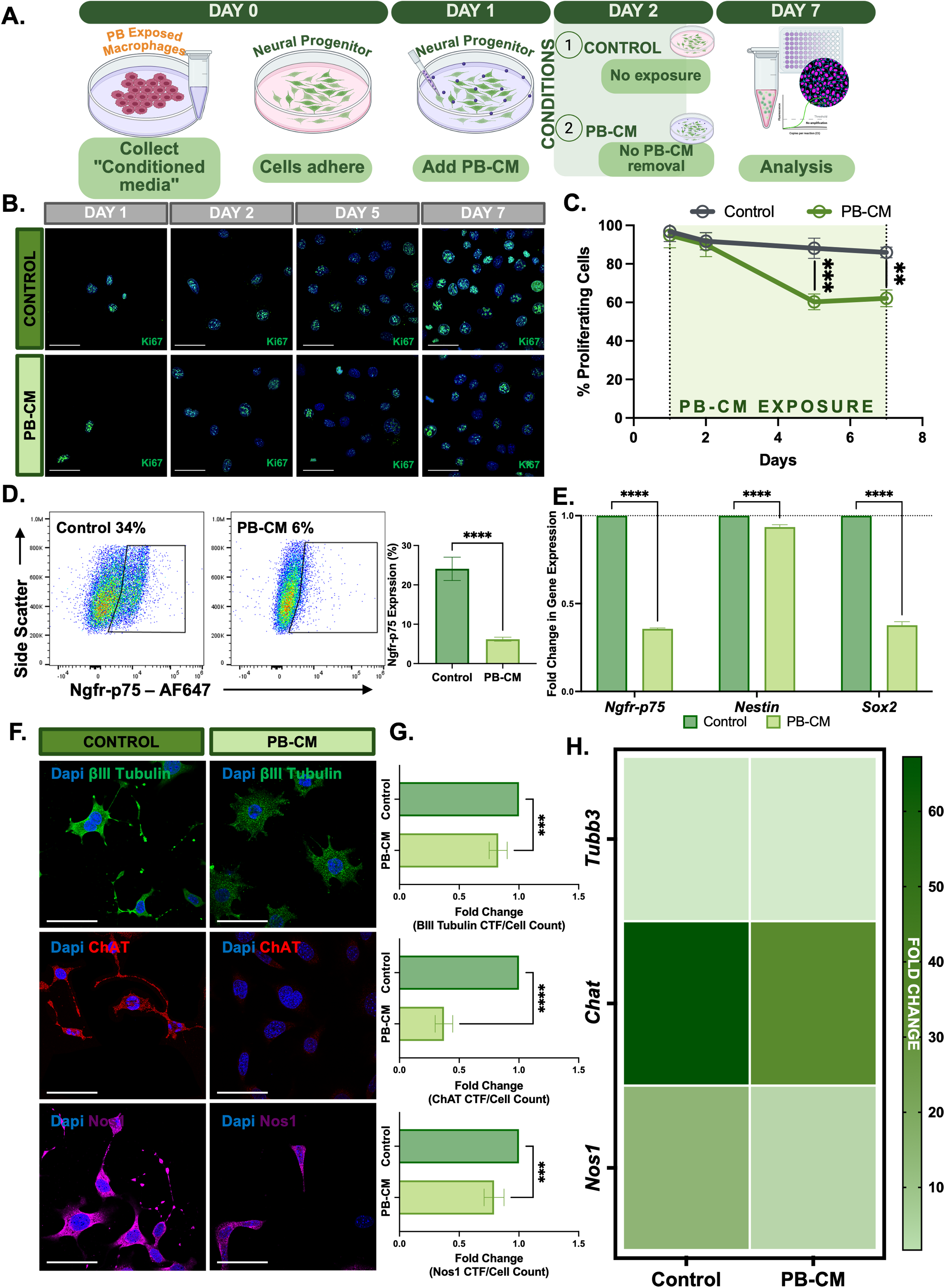
Inflammatory secretome from PB-exposed macrophages disrupts IM-FEN proliferation and neurogenic potential. (A) Schematic of experimental timeline for treating IM-FENs with conditioned media collected from PB-exposed macrophages (PB-CM). Macrophages were exposed to PB for 24hrs, and conditioned media was collected immediately following exposure and stored (PB-CM). Enteric neuronal progenitor cells were adhered to culture surfaces, and treated with PB-CM prior to differentiation on Day 1. Cultures of IM-FENs were maintained in undifferentiated progenitor states to evaluate proliferation and stem cell identity. Cultures of IM-FENs were differentiated beginning at Day 2, for 5 days leading up to Day 7 in the presence or absence of PB-CM, to analyze their differentiation capability. (B) Representative immunofluorescence images showing Ki67 (green), a marker of proliferating cells, in untreated and PB-CM treated IM-FEN cultures at the indicated timepoints. Nuclei are counterstained with DAPI (blue). Scale bars = 15 µm. (C) Quantification of Ki67⁺ proliferating cells over 7 days. Data are presented as mean ± SEM. ****p<0.0001, **p<0.01, two-way ANOVA; n>5. (D) Flow cytometry analysis of neural progenitor marker Ngfr-p75 expression in control and PB-CM treated IM-FENs. Data are presented as mean ± SEM. ****p<0.0001, unpaired t-test; n>9. (E) Relative gene expression of neural stem markers *(Ngfr-p75*, *Nestin*, *Sox2)* in PB-CM treated IM-FENs, normalized to untreated controls (dotted line = 1). Data are presented as mean ± SEM. ****p<0.0001, ‘ns’ = not significant; two-way ANOVA; n>4. (F, G) Untreated or PB-CM treated IM-FENs were differentiated into several neuronal subtypes. (F) Representative immunofluorescence images showing expression of neuronal markers: βIII-tubulin (green), ChAT (red), and Nos1 (magenta) in control and PB-CM treated differentiated IM-FENs. Nuclei are counterstained with DAPI (blue). Scale bars = 20 µm. (G) Quantification of differentiated neuronal populations expressing βIII-tubulin, ChAT, and Nos1. Data are presented as mean ± SEM. ****p<0.0001, ***p<0.001; unpaired t-test; n>5. (H) Heatmap displaying relative gene expression fold changes of *Tubb3*, *Chat*, and *Nos1* in untreated and PB-CM treated differentiated IM-FENs, normalized to undifferentiated controls.

#### 3.8.1 PB-CM reduces enteric neuronal progenitor proliferation and depletes progenitor identity

Ki67 immunostaining revealed a marked reduction in proliferative IM-FENs following continuous exposure to PB-CM (**Figure 7B**), compared to untreated controls. Quantitative analysis showed a 31% decrease in Ki67^+^ cells at day 5, and 28% at day 7 in PB-CM treated cultures relative to untreated controls (**\*\*p<0.01; Figure 7C**), suggesting a potential impairment in self-renewal ability. This reduced proliferative capacity was accompanied by an 82% decline in Ngfr-p75^+^ cells (a marker of stem-like identity) (**\*\*\*\*p<0.0001; Figure 7D**), indicating that prolonged exposure to inflammatory CM severely compromises neural progenitor maintenance. The gating strategy for all flow analysis is provided in **Supplementary Figure 7.** Gene expression analysis supported these findings, showing significant downregulation of neural stem cell markers *Ngfr-p75 (***0.36-fold**)*, Nestin (***0.94-fold**), and *Sox2 (***0.38-fold**) in PB-CM treated IM-FENs (**\*\*\*\*p<0.0001; Figure 7E**), consistent with loss of stem-like phenotype.

#### 3.8.2 Inflammatory PB-CM impairs neuronal differentiation of IM-FENs

Neural progenitor cells are essential for maintaining productive neuroplasticity in response to inflammation. To understand how inflammation affects this regenerative potential in neuronal progenitor cells, we evaluated neuronal differentiation in the presence of PB-CM. When stimulated to differentiate with exogenously added factors, neuronal progenitor cells demonstrated an impaired ability to differentiate with reduction in βIII-tubulin, ChAT, and Nos1 in PB-CM treated cultures compared to controls (**Figure 7F, G**). This loss of neuronal markers was supported by gene expression analysis, which revealed 49% and 52% reductions in *Chat* and *Nos1* expression (**\*\*\*p<0.001; Figure 7H**). These results reveal a marked impairment in neurogenesis under PB-mediated inflammation, characterized by loss of neuronal identity and differentiation capacity.

These findings support our hypothesis that PB-induced, macrophage-mediated inflammation impairs enteric neural repair and contributes to dysmotility. In PB-exposed bioengineered colon assembloids, we observed sustained inflammation, and activation of cell death pathways, as evidenced by transcriptomic profiling (**Figures 3**) and elevated levels of inflammatory cytokines (**Figure 4**), alongside major neuronal losses and inflammatory macrophage accumulation (**Figure 5**). Collectively, this suggests that PB exposure initiates a strong macrophage-driven inflammatory response that persists beyond the initial insult, leading to long-term disruption of neurogenesis and neural circuit integrity. This chronic neuroinflammation (despite PB removal) likely explains the loss of functional motility observed in the PB-exposed bioengineered colon assembloid.

## CONCLUSIONS

Through our work, we have demonstrated the fabrication of a multi-cellular bioengineered colon assembloid, that structurally and functionally integrates key cell types of the colonic muscularis externa: smooth muscle cells, macrophages and an enteric neural circuit. The platform enables real-time quantitative assessment of colonic contractility, directly linking neuro-immune interactions to functional contractility outputs. Our results demonstrate the bioengineered colon assembloid is sensitive to inflammatory perturbations, inducing measurable changes in macrophage activation, enteric neuronal plasticity, and ultimately, colonic contractility. The platform also provides a tightly controlled environment for isolating cellular variables to interrogate molecular mechanisms in the future to investigate how neuro-inflammation contributes to colonic dysmotility in FGIDs.

## ETHICAL CONSIDERATIONS

This work was conducted in a laboratory approved by the Institutional Biosafety Committee at Texas A&M University. No animal use or institutional review board approval was required, since there is no human or vertebrate animal research.

## FUNDING

This work was supported by the Department of Defense Gulf War Illness Research Program W81XWH-21-0477 (S.A.R.), Department of Defense Toxic Exposures Research Program HT9425-24-1-0998 (S.A.R.), Texas A&M University Avilés-Johnson Graduate Fellowship Program (C.A.C.), the National Defense Science and Engineering Graduate (NDSEG) Fellowship Program (C.A.C.) through the Army Research Office, and NIH R01 DK080684 (S.S.).

## DATA TRANSPARENCY STATEMENT

Data supporting the findings of this study will be available from the corresponding author upon written request. RNA-Seq datasets are deposited on the NCBI Gene Expression Omnibus database, under ID: GSE316347.

## REPORTING GUIDELINES

The data reported in this manuscript is not related to randomized trials, case reports or animal studies. Therefore, reporting guidelines are not applicable.

## CONFLICTS OF INTEREST

The authors report no conflicting conflicts for interests.

## AUTHOR CONTRIBUTIONS

Conceptualization: S.A.R.;SS Data curation: C.A.C., K.O.S; Formal analysis: C.A.C.,K.O.S, S.A.R.; Funding acquisition: S.A.R.; S.S; Investigation: C.A.C.,K.O.S, A.S.; Methodology: C.A.C., K.O.S, S.A.R.; Project administration: S.A.R.; Resources: S.A.R., S.S.; Software: C.A.C.; Supervision: S.A.R.; Validation: C.A.C., S.A.R.; Visualization: C.A.C., K.O.S, A.S., A.V.,A.A.M, S.A.R.; Writing – original draft: C.A.C., S.A.R. ; Writing – review & editing: S.A.R, S.S.

**PREPRINT**: https://doi.org/10.1101/2025.08.07.669171

## LIST OF ABBREVIATIONS

α-SMA: alpha smooth muscle actin
β-NGF: beta nerve growth factor
ACh: acetylcholine
BMP-2: bone morphogenetic protein-2
CCL2, CCL11: C-C-motif chemokine ligand 2/11
ChAT: Choline acetyltransferase
DAPI-4,6: diamidino-2-phenylindole
ENS: enteric nervous system
FGID: functional gastrointestinal disorders
GWI: Gulf War Illness
iBMDM: immortalized murine bone marrow derived macrophages
IBD: inflammatory bowel disease
iCSMC: immortalized colonic smooth muscle cells
IM-FEN: immorto mouse fetal enteric neuron
IL-1β,6,10: interleukin
NGFR: nerve growth factor receptor
nNOS/NOS1: neuronal nitric oxide synthase
PB: pyridostigmine bromide
TIMP-1: tissue inhinitor of metalloproteinase
TNF-α: tumor necrosis factor alpha
TTX: tetrodotoxin

## Notes

### Competing Interest Statement

The authors have declared no competing interest.

### Summary of Updates

Updated to match accepted manuscript in revised form

