## Supplementary Information for "Macrophage Immune-Competent Colon Assembloids for Functional Interrogation of Neuroinflammation-Induced Colonic Dysmotility"

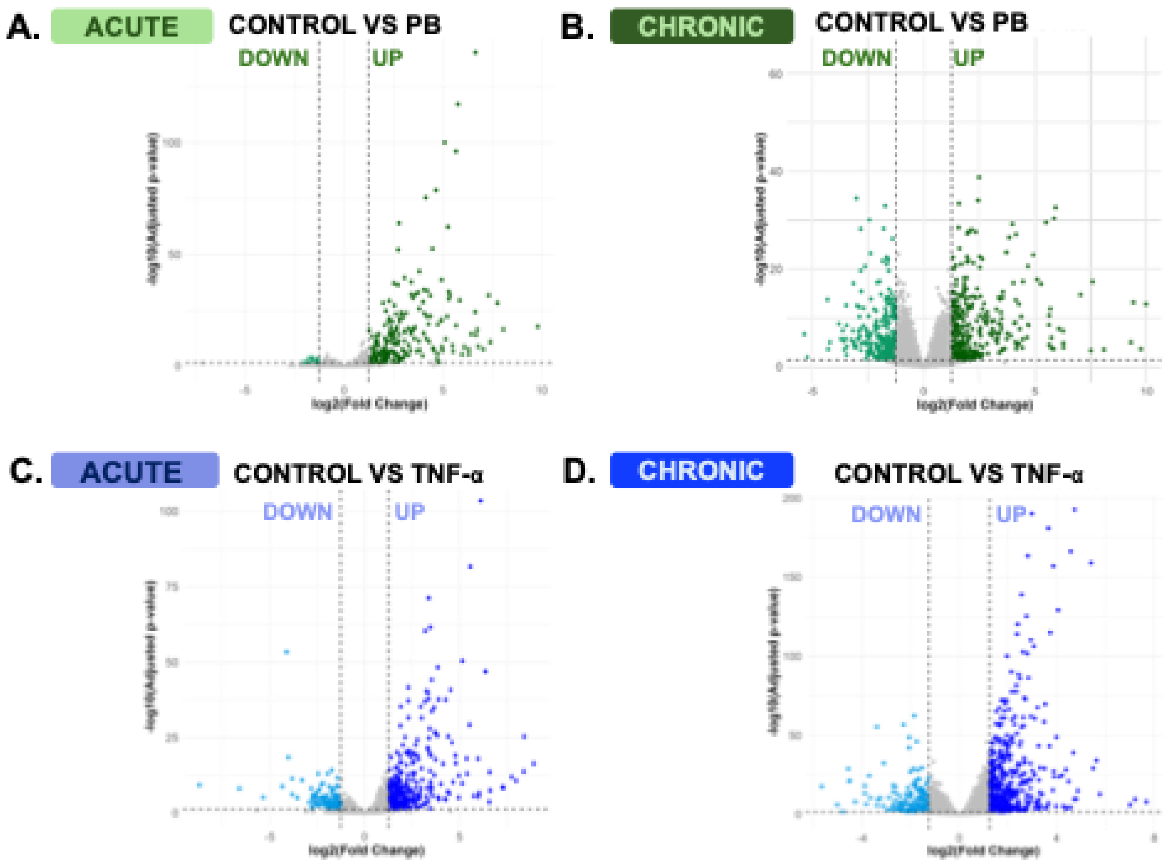


**Supplementary Figure 1.** *Differential Gene Expression Profiles in Acute and Chronic Recovery from PB or TNF-α Exposure.* (A, B) Volcano plots illustrating differentially expressed genes (DEGs) in bioengineered colon assembloids following acute (A) and chronic (B) exposure to PB compared to untreated controls. (C, D) Volcano plots illustrating differentially expressed genes (DEGs) in bioengineered colon rings following acute (C) and chronic (D) exposure to TNF-α compared to untreated controls. Upregulated and downregulated genes are indicated as log2FC > ± 1.25. Colored points denote significantly upregulated (right) or downregulated (left) genes (green = PB, blue = TNF-α), while nonsignificant genes are shown in grey.

PB exposure resulted in upregulation of *Cd74*[101], *Cxcl9*[102], *Ciita*[103, 104], *Cx3cl1*[105], *Ccl2*[106]. This signature is strongly associated with activating innate immune responses to inflammation. PB exposure was also associated with a significant down regulation of *Adam19*[107], *Fyn*[108], *Nptxr*[109], and *Dkk2*[110]. This signature is strongly associated with impaired neural development and structural tissue organization (**Supplementary Figure 1A**). In chronic conditions, lingering effects from one-time PB exposure continued to drive strong immunologic signatures, with upregulation of *Casp1*[111], *Ccl11*[40], *Stat1*[112], and downregulation of *Nes*[18], *Npy*[113], *and Nell2*[114]. This gene signature reflects chronic neuroinflammation and impaired neurogenesis, supporting that PB-induced inflammation drives lasting neuronal damage in the bioengineered colon assembloid (**Supplementary Figure 1B**).

Acute TNF-α exposure also induced gene signatures associated with inflammation, albeit through a slightly different set of genes (upregulation of *Nes, Relb*[55], *Mmp9*[115], *Sdc4*[116], *Csf1*[117], and *Wisp1*[118]). A similar disruption in neurogenesis was observed with TNF-α treatment as well (downregulation of *Eya4, Sema6a, Mef2c, Sox2*[119], and *Nrxn2*[120, 121] (**Supplementary Figure 1C**) (log2FC > ±1.25 for all). Chronic effects from one-time TNF-α exposure elicited gene expression changes indicative of extracellular matrix (ECM) remodeling, including upregulation of *Col7a1*[122] and *Mmp9*[123]. Concurrently, downregulation of *Cd40*[124], *Ciita,* and *Csf2* suggests a shift away from immune activation (**Supplementary Figure 1D**). Further analysis of in-depth differences between PB and TNF-α can be found in **Supplementary Figure 2**.

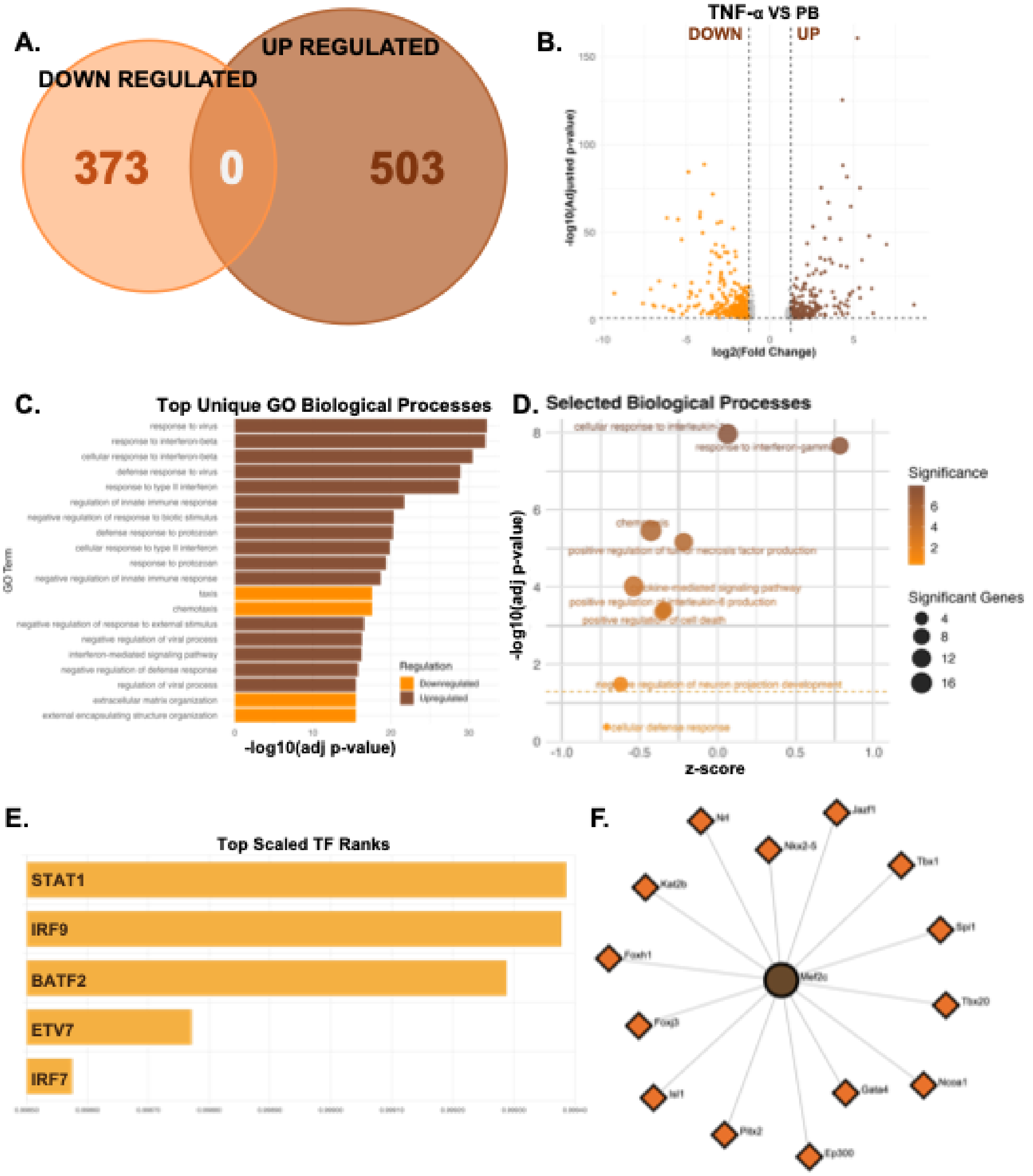


**Supplementary Figure 2.** *Head-to-head comparison of PB and TNF-α reveals unique inflammatory signatures.* (A) Venn diagram comparing significantly upregulated and downregulated genes in PB-exposed colon assembloids relative to TNF-α exposed assembloids. (B) Volcano plot showing differentially expressed genes (DEGs) in PB-treated bioengineered colon assembloids compared to TNF-α-treated rings. Upregulated and downregulated genes are indicated as log2FC > ± 1.25. (C) Top unbiased Gene Ontology (GO) Biological Process (BP) terms significantly enriched in PB-treated assembloids relative to TNF-α treatment. (D) Selected GO BP terms plotted by z-score and –log10(adjusted p-value), highlighting relative pathway enrichment and statistical

significance. Z-scores reflect the deviation in the proportion of significant genes per pathway compared to the average, with positive values indicating higher-than-expected enrichment. Dot size denotes the number of significant genes per term, and color intensity corresponds to adjusted p-value significance. (E) Top-ranked predicted transcription factors associated with upregulated gene expression in PB-treated rings. (F) Transcription factor prediction was performed using the ChEA3 platform (https://amp.pharm.mssm.edu/ChEA3)[45], using the "TopRank" integrated library and inputting all upregulated DEGs for each condition. Network visualization was carried out using NetworkAnalyst (https://www.networkanalyst.ca)[46] by submitting the top five upregulated genes per condition to generate protein–protein interaction networks.
Inferred transcriptional regulatory network for PB-treated samples, displaying predicted interactions between transcription factors (diamonds) and their target genes (circles).

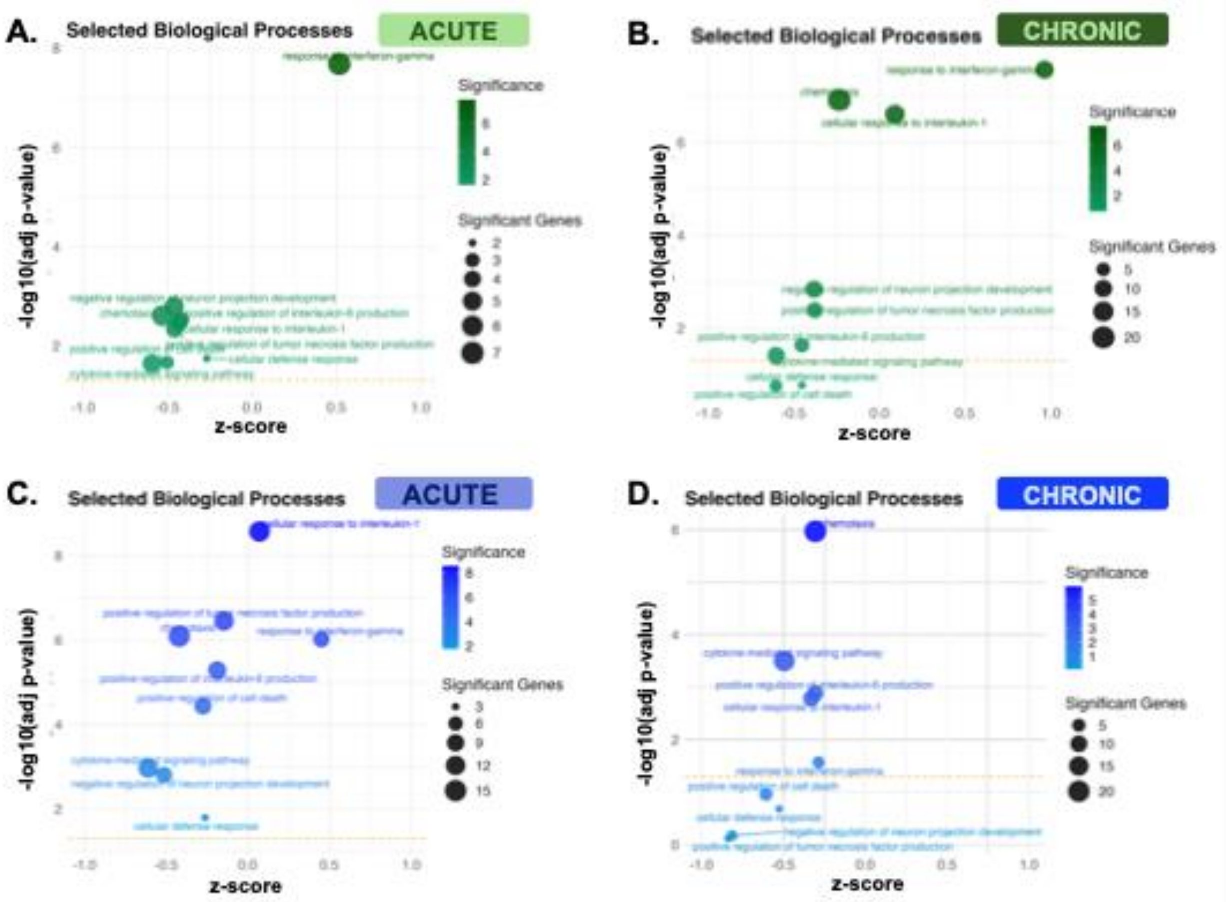


In addition to the full GO output, a focused set of terms relevant to immune signaling, apoptosis, and neuroinflammation was selected for targeted enrichment: GO:0071347, GO:0019221, GO:0032755, GO:0032760, GO:0006935, GO:0006968, GO:0010942, GO:0034341, GO:0010977.

**Supplementary Figure 3.** *Gene Ontology Biological Process Enrichment Analysis in PB and TNF-α Bioengineered Colon Assembloids.* (A, B) Selected Gene Ontology (GO) Biological Process (BP) terms for acute effects (A) and chronic effects (B) from PB exposure.

(C, D) Selected GO BP terms for acute (C) and chronic (D) effects from TNF-α exposure. Pathways are plotted by z-score and –log10(adjusted p-value), highlighting pathway-level enrichment. The z-score reflects the deviation of the proportion of significant genes in each pathway from the average across all pathways, with positive values indicating higher-than-average enrichment. Circle size corresponds to the number of significant genes, and darker color indicates higher statistical significance. Node color distinguishes condition type (green, PB; blue, TNF-α).

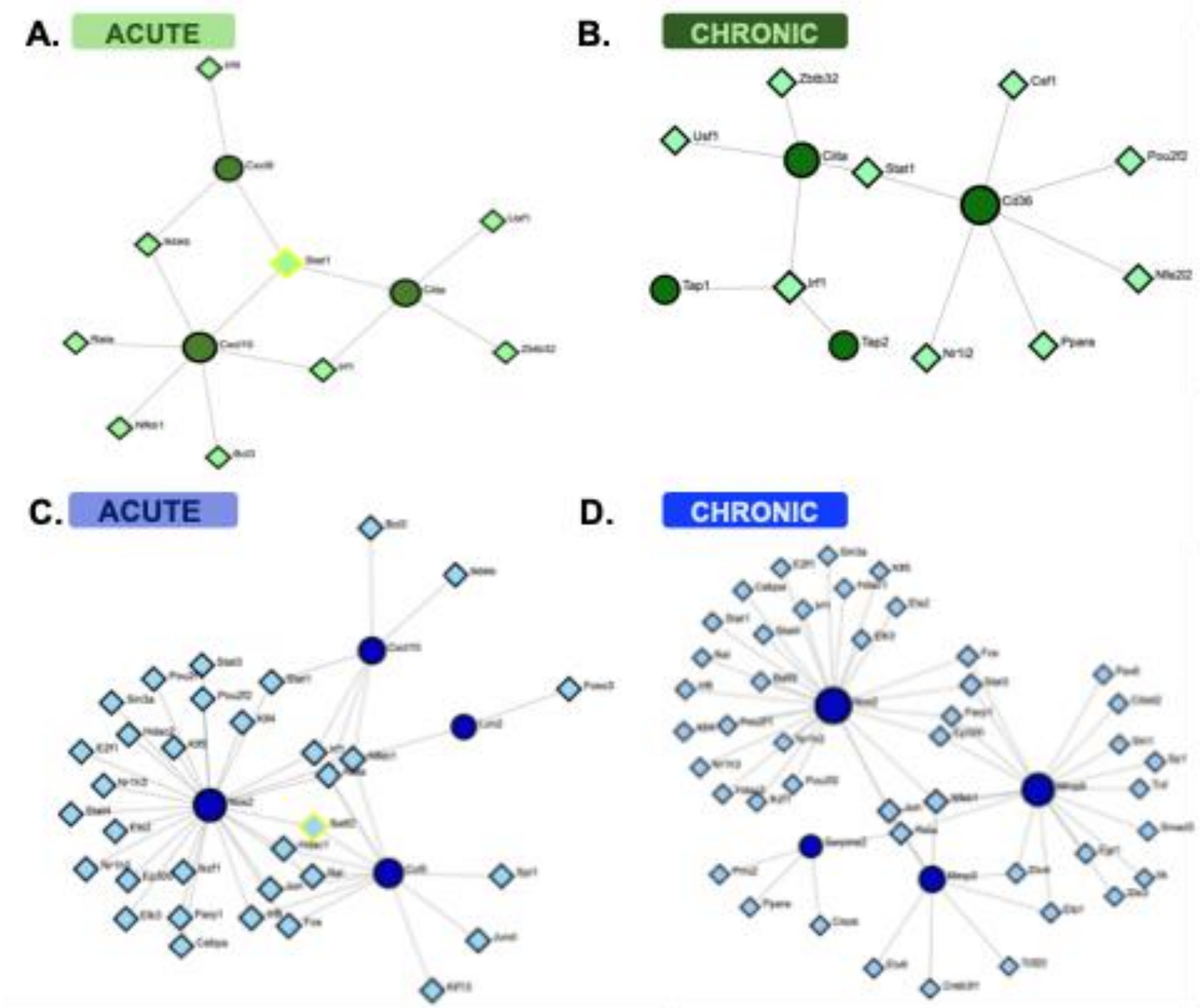


**Supplementary Figure 4.** *Transcription factor–gene regulatory networks altered by PB or TNF-α exposure.* Transcription factor prediction was performed using the ChEA3 platform (https://amp.pharm.mssm.edu/ChEA3)[45], using the "TopRank" integrated library and inputting all upregulated DEGs for each condition. Network visualization was carried out using NetworkAnalyst (https://www.networkanalyst.ca)[46] by submitting the top five upregulated genes per condition to generate protein–protein interaction networks.
(A, B) Predicted transcriptional regulatory networks for acute (A) and chronic (B) PB exposure. (C, D) Predicted transcriptional regulatory networks for acute (C) and chronic (D) TNF-α exposure. Diamonds denote transcription factors and circles represent target genes, with edges indicating inferred regulatory relationships. Node color indicates exposure type (green, PB; blue, TNF-α), and node size reflects network connectivity (degree of interactions). Yellow-highlighted nodes represent top-ranked hub regulators identified in the analysis.

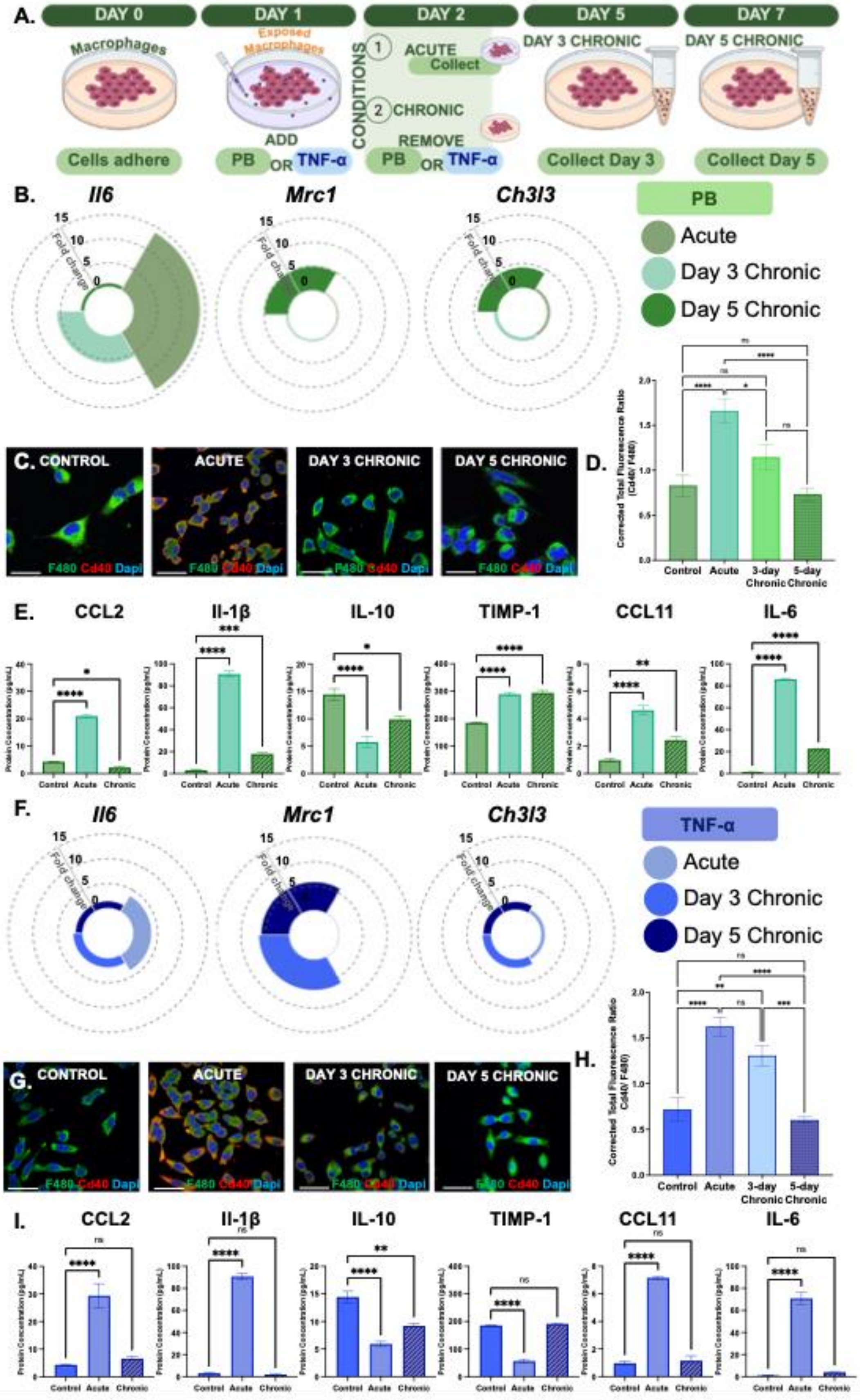
A.
DAY 0
DAY 1
DAY 2
DAY 5
DAY 7
Macrophages
Exposed Macrophages
CONDITIONS
1 ACUTE
Collect
2 CHRONIC
REMOVE
DAY 3 CHRONIC
DAY 5 CHRONIC
Cells adhere
ADD
PB OR TNF-α
PB OR TNF-α
Collect Day 3
Collect Day 5
B.
Il6
Mrc1
Ch3l3
Fold change
PB
Acute
Day 3 Chronic
Day 5 Chronic
C. CONTROL
ACUTE
DAY 3 CHRONIC
DAY 5 CHRONIC
F480 Cd40 Dapi
D.
Corrected Total Fluorescence Ratio (Cd40/ F480)
Control
Acute
3-day Chronic
5-day Chronic
E.
CCL2
Il-1β
IL-10
TIMP-1
CCL11
IL-6
Protein Concentration (pg/mL)
Control
Acute
Chronic
F.
Il6
Mrc1
Ch3l3
TNF-α
Acute
Day 3 Chronic
Day 5 Chronic
G. CONTROL
ACUTE
DAY 3 CHRONIC
DAY 5 CHRONIC
H.
I.
CCL2
Il-1β
IL-10
TIMP-1
CCL11
IL-6

**Supplementary Figure 5.** *Time-Dependent Resolution of Macrophage Activation and Secretory Profiles Following Acute PB and TNF-α Exposure* (A) Schematic outlining the experimental timeline for macrophage stimulation (one-time exposure) and recovery. Immortalized bone marrow-derived macrophages (iBMDMs) were treated with 100 µM PB or 100 ng/mL TNF-α for 24 hours (acute), followed by either immediate sample collection or a recovery period in fresh media for 3 or 5 days and evaluation of lingering inflammation (chronic). (B) Circular bar plots showing average gene expression fold changes of macrophage markers (*Il6*, *Mrc1*, *Ch3l3*) following acute and chronic PB exposure. (C) Representative immunofluorescence images of macrophages stained for F4/80 (green), Cd40 (red), and DAPI (blue) under control, acute, and chronic PB conditions. Scale bar = 15µm. (D) Quantification of Cd40 expression in control and PB-treated macrophages. Data are presented as mean ± SEM. ****p<0.0001, *p<0.05, 'ns' = not significant; ordinary one-way ANOVA; n>5. (E) Cytokine levels following PB treatment in macrophages. Data are presented as mean ± SEM. ****p<0.0001, ***p<0.001, **p<0.01, *p<0.05; ordinary one-way ANOVA; n>3. (F) Average gene expression fold changes of macrophage markers (*Il6, Mrc1, Ch3l3*) following TNF-α treatment. (G) Representative immunofluorescence images of macrophages stained for F4/80 (green), Cd40 (red), and DAPI (blue) under control, acute, and chronic TNF-α conditions. Scale bar = 15µm. (H) Quantification of Cd40 expression in control and TNF-α treated macrophages. Data are presented as mean ± SEM. ****p<0.0001, ***p<0.001, **p<0.01, 'ns' = not significant; ordinary one-way ANOVA; n>5. (E) Cytokine levels following TNF-α treatment in macrophages. Data are presented as mean ± SEM. ****p<0.0001, **p<0.01, 'ns' = not significant; ordinary one-way ANOVA; n>3.

Acute PB exposure significantly upregulated pro-inflammatory gene expression of *Il6* by 14-fold, confirming activation of a classical inflammatory response (****p<0.0001; Supplementary Figure 5B**). Upon removal of PB, *Il6* gene expression decreased in a time-dependent manner declining substantially over 7 days, with a steady and concurrent upregulation of *Ch3l3* and *Mrc1*. In parallel, inflammatory Cd40 expression peaked with acute PB exposure (2-fold increase; ****p<0.0001; **Supplementary Figure 5C, D**) with a gradual decline over 7 days. Secreted cytokine patterns also mirrored the acute inflammation (increases in CCL2, IL-1β, CCL11 and IL-6) with an active decline towards baseline over time (**Supplementary Figure 5E**). Together, these results demonstrated that PB induced inflammation in macrophages was robust, but reversible, with a shift towards resolution of inflammation in the absence of other cell types amplifying tissue damage. Acute TNF-α exposure significantly upregulated *Il6* gene expression by 3.8-fold (****p<0.0001; Supplementary Figure 5F**), confirming induction of a pro-inflammatory macrophage activation. Similar to PB exposure, *Il6* expression declined progressively to near-baseline levels (1.5-fold; ****p<0.0001**), with increased gene expression of reparative markers, *Mrc1* and *Ch3l3* (****p<0.0001; Supplementary Figure 5F**). Macrophages also transiently increased Cd40 expression, with a gradual decline following removal of TNF- α and recovery (**Supplementary Figure 5G, H**). Secreted cytokines also matched these activation patterns (**Supplementary Figure 5I**). This temporal pattern parallels the response to PB, highlighting the ability of macrophages to consistently resolve inflammation across different inflammatory contexts.

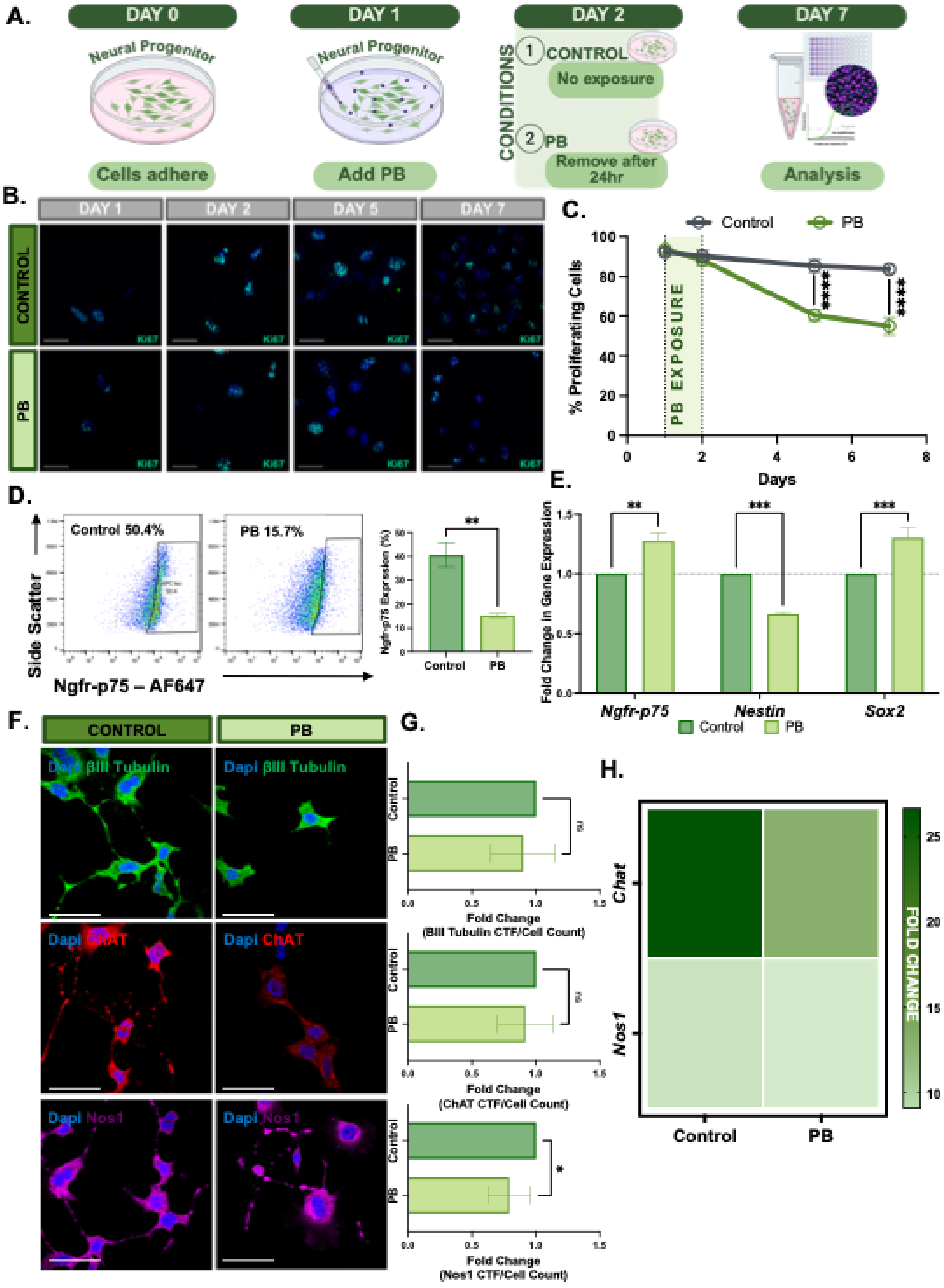

A.
DAY 0
Neural Progenitor
Cells adhere
DAY 1
Neural Progenitor
Add PB
DAY 2
CONDITIONS
1 CONTROL
No exposure
2 PB
Remove after 24hr
DAY 7
Analysis
B.
DAY 1
DAY 2
DAY 5
DAY 7
CONTROL
PB
Ki67
C.
Control
PB
% Proliferating Cells
PB EXPOSURE
****
****
Days
D.
Control 50.4%
PB 15.7%
Side Scatter
Ngfr-p75 – AF647
**
Ngfr-p75 Expression (%)
Control
PB
E.
Fold Change in Gene Expression
**
***
***
Ngfr-p75
Nestin
Sox2
Control
PB
F.
CONTROL
PB
Dapi βIII Tubulin
Dapi ChAT
Dapi Nos1
G.
Control
PB
ns
Fold Change (BIII Tubulin CTF/Cell Count)
Fold Change (ChAT CTF/Cell Count)
*
Fold Change (Nos1 CTF/Cell Count)
H.
Chat
Nos1
Control
PB
FOLD CHANGE

**Supplementary Figure 6.** *PB exposure alone does not impair IM-FEN fate and neuronal differentiation.* (A) Schematic of the experimental timeline for IM-FEN treatment with pyridostigmine bromide (PB). (B) Representative immunofluorescence images of Ki67 (green) in control and PB-treated IM-FENs at indicated time points, assessing cell proliferation. Nuclei are counterstained with DAPI (blue). Scale bars = 15 µm. (C) Quantification of the percentage of $Ki67^+$ proliferating cells across 7 days. Data are presented as mean ± SEM. ****p<0.0001, two-way ANOVA; n>5. (D) Flow cytometry analysis of the neural progenitor marker Ngfr-p75 in control and PB-treated IM-FENs. (E) Relative gene expression of neural stem markers *(Ngfr-p75*, *Nestin*, *Sox2)* in PB-treated IM-FENs, normalized to untreated controls (dotted line = 1). Data are presented as mean ± SEM. ***p<0.001, **p<0.01; two-way ANOVA; n>4. (F, G) Untreated or PB-treated IM-FENs were differentiated into several neuronal subtypes. (F) Representative immunofluorescence images showing expression of neuronal markers: βIII-tubulin (green), ChAT (red), and Nos1 (magenta) in control and PB-treated differentiated IM-FENs. Nuclei are counterstained with DAPI (blue). Scale bars = 20 µm. (G) Quantification of differentiated neuronal populations expressing βIII-tubulin, ChAT, and Nos1. Data are presented as mean ± SEM. *p<0.05; 'ns' = not significant; unpaired t-test; n>5. (H) Heatmap displaying relative gene expression fold changes of *Chat* and *Nos1* in untreated and PB-treated differentiated IM-FENs, normalized to undifferentiated controls.

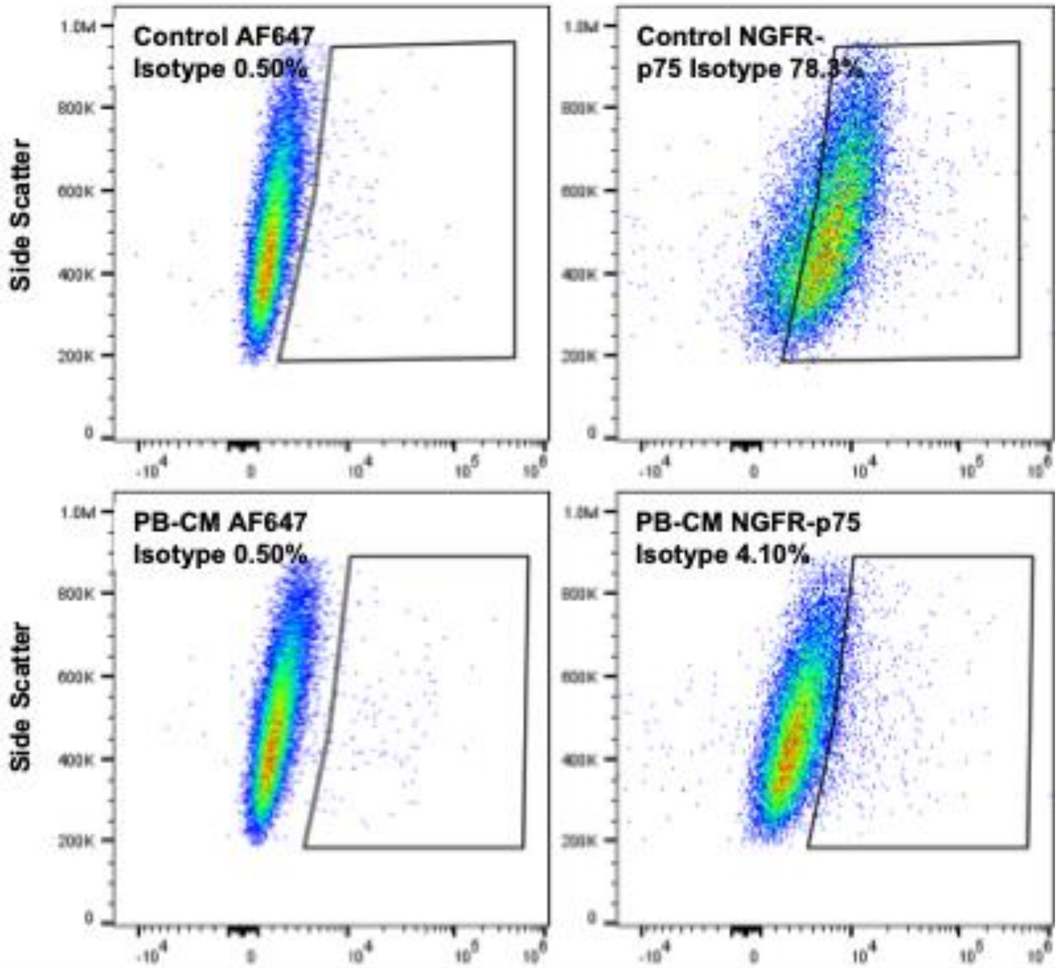


**Supplementary Figure 7.** *NGFR-p75 flow cytometry gating technique using isotype controls.* (A) Polygon gate (outlined in black) was drawn on FlowJo to create a 0.5% background cut off gate in the isotype control graph (left). The same gate was applied to the antibody-stained graph (right) to establish the percentage of the population expressing NGFR-p75.

**Supplementary Table 1**

**Supplementary Table 1**. List of antibodies used in immunohistochemistry.

| Antibody | Conjugation | Manufacturer | Catalog # | RRID |
|---|---|---|---|---|
| F4/80 | Alexa Fluor 488 | Abcam | ab204266 | AB_2943479 |
| Ym1 | PE | Abcam | ab211621 | AB_3083045 |
| Cd40 | Alexa Fluor 647 | Abcam | ab275158 | AB_2839768 |
| Ngfr-p75 | Alexa Fluor 647 | Santa Cruz Biotechnology | sc-271708 | AB_10714958 |
| βIII-Tubulin | Alexa Fluor 488 | invitrogen | 53-4510-82 | AB_1107000 |

| | | | | |
|---|---|---|---|---|
| ChAT | FITC; Alexa Fluor 647 | Santa Cruz Biotechnology | sc-55557 | AB_2291743 |
| Nos1 | Alexa Fluor 647 | Santa Cruz Biotechnology | sc-5302 | AB_631879 |
| SMα | Coralite 594 | Proteintech | cl595-14395 | AB_2889885 |
| Ki67 | Alexa Fluor 488 | Cell signaling Technologies | 11882S | AB_2797703 |

**Supplementary Table 2**

**Supplementary Table 2.** List of primers used in RT-qPCR analysis of gene expression.

| Mouse Primer | Gene Sequences |
|---|---|
| *Gapdh* | F: AGGTCGGTGTGAACGGATTTG<br>R: GGGGTCGTTGATGGCAACA |
| *Nos1* | F: CCCAACGTCATTTCTGTCCGT<br>R: TCTACCAGGGGCCGATCATT |
| *Chat* | F: GGCCATTGTGAAGCGGTTTG<br>R: GCCAGGCGGTTGTTTAGATACA |
| *Tubb3* | F: CCCAGCGGCAACTATGTAGG<br>R: CCAGACCGAACACTGTCCA |
| *Ngfr-p75* | F: TGCCGATGCTCCTATGGCTA<br>R: CTGGGCACTCTTCACACACTG |
| *Sox2* | F: GCGGAGTGGAAACTTTTGTCC<br>R: GGGAAGCGTGTACTTATCCTTCT |
| *Nes* | F: CCCACCTATGTCTGAGGCTC<br>R: GGGCTAAGGAGGTTGGATCAT |
| *Mrc1* | F: CTCTGTTCAGCTATTGGACGC<br>R: TGGCACTCCCAAACATAATTTGA |
| *Ch3l3* | F: CAG GTC TGG CAA TTC TTC TGA A<br>R: GTC TTG CTC ATG TGT GTA AGT GA |
| *Il6* | F: TCT ATA CCA CTT CAC AAG TCG GA<br>R: GAA TTG CCA TTG CAC AAC TCT TT |
| *Cald1* | F: AGA AGG AGT TTG ATC CGA CCA<br>R: CAT TTT CGG CGG AGT CAT TTT G |
| *Tagln* | F: ACC AAA AAC GAT GGA AAC TAC CG<br>R: GTG AAG TCC CTC TTA TGC TCC T |
| *Acta2* | F: GGC ACC ACT GAA CCC TAA GG<br>R: ACA ATA CCA GTT GTA CGT CCA GA |
